# Endocytic control of cell-autonomous and non-cell-autonomous functions of p53

**DOI:** 10.1101/2025.08.16.670648

**Authors:** Roberta Cacciatore, Andrea Basile, Stefano Freddi, Irene Schiano Lomoriello, Carlo Ribelle Zucca, Giuseppe Ciossani, Luigi Scietti, Alessandro Cuomo, Simona Ronzoni, Simone Pelicci, Mario Faretta, Elena Zaccheroni, Giuliana Pelicci, Vittoria Matafora, Angela Bachi, Rosalind Helen Gunby, Salvatore Pece, Sara Sigismund, Letizia Lanzetti, Ivan Nicola Colaluca, Pier Paolo Di Fiore

**Author notes:** Corresponding author: Pier Paolo Di Fiore. Equal first authors. Equal last authors.

## Abstract

NUMB is an endocytic protein with tumor suppressor activity, largely mediated by its ability to inhibit p53 degradation. This function depends on the inclusion of a short alternatively spliced exon (Ex3) in NUMB, although the mechanistic link between endocytosis and p53 regulation remains unclear. Here, we show that the Ex3-encoded sequence directs NUMB to the plasma membrane, where it forms a complex with the endocytic adaptor SNX9. This complex recruits p53 in a SNX9-dependent manner and is internalized and trafficked to multivesicular bodies, culminating in exosomal secretion, in a process requiring both SNX9 and NUMB. Exosomal p53 is taken up by recipient cells and translocated to the nucleus, where it activates p53-dependent transcriptional and phenotypic programs. These findings suggest that exosome-mediated p53 transfer may contribute to the establishment of a tumor-suppressive microenvironment.

## 1. INTRODUCTION

NUMB is a tumor suppressor gene implicated in the pathogenesis of various cancers, including those of the breast, prostate, lung, stomach, and liver ^1–18^. Initially discovered in *Drosophila melanogaster* as a key regulator of cell fate, NUMB orchestrates the asymmetric division of cells during mitosis, thereby determining the fate of daughter cells in a cell-autonomous manner ^19–22^. In *Drosophila*, NUMB’s primary role is to antagonize the signaling activity of the NOTCH receptor through its well-established function as an endocytic regulator ^23–31^. This mechanism is evolutionarily conserved ^32–35^; however, in chordates, including mammals, the emergence of an alternatively spliced exon (Exon 3, Ex3) ^27^ adds further complexity to the regulation of cellular fates by NUMB. Specifically, the Ex3-encoded sequence allows NUMB to interact with MDM2, the principal E3 ligase responsible for the ubiquitination and ensuing proteasomal degradation of the tumor suppressor p53 ^1, 2, 27^. Consequently, NUMB stabilizes p53, a function critical for the maintenance of stem cell (SC) homeostasis across various tissues, including the mammary gland ^5, 8, 13^. Notably, NUMB ablation disrupts SC division, shifting it from an asymmetric pattern (1 SC → 1 SC + 1 progenitor) to a symmetric one (1 SC → 2 SCs), resulting in an expansion of the SC pool ^5, 8, 13^, a characteristic of cancer stem cells (CSCs). Thus, NUMB plays a pivotal physiological role by suppressing signaling through NOTCH, a proto-oncogene, and sustaining the activity of p53, a key tumor suppressor.

Consistent with its physiological role, NUMB loss-of-function (LOF) contributes significantly to oncogenesis, as exemplified in breast cancer, where NUMB alterations are linked to more aggressive disease progression. In breast cancer, NUMB LOF occurs in approximately 65% of cases ^36^, arising through several mechanisms: i) reduction of NUMB protein levels due to enhanced degradation mediated by the CRL7^FBXW8^ E3 ligase complex ^36^, ii) splicing defects that reduce the expression of Ex3-containing isoforms ^1^, and iii) aberrant phosphorylation events that impair NUMB’s partitioning during cell division ^5^.

In mammals, NUMB exists in four major isoforms, generated by the alternative splicing of two exons, Ex3 and Ex9 (see Fig. 1A) ^37^. Ex3 encodes a short stretch of 11 amino acids within the phosphotyrosine-binding (PTB) domain at the N-terminus of the protein, resulting in two distinct PTB isoforms: one containing Ex3 and one lacking it. The only known function of the Ex3-coded sequence is to bind MDM2, thereby stabilizing p53. The relevance of this function to cancer is underscored by the observation that approximately 18% of breast cancers exhibit reduced expression of Ex3-containing NUMB isoforms ^36^, an alteration that correlates with an aggressive disease course and poor prognosis, independently of other risk factors ^1^. Furthermore, breast cancers with reduced levels of Ex3-containing isoforms exhibit heightened chemoresistance, a phenomenon that could be mechanistically linked to decreased p53 levels^1^.

**Figure 1.**
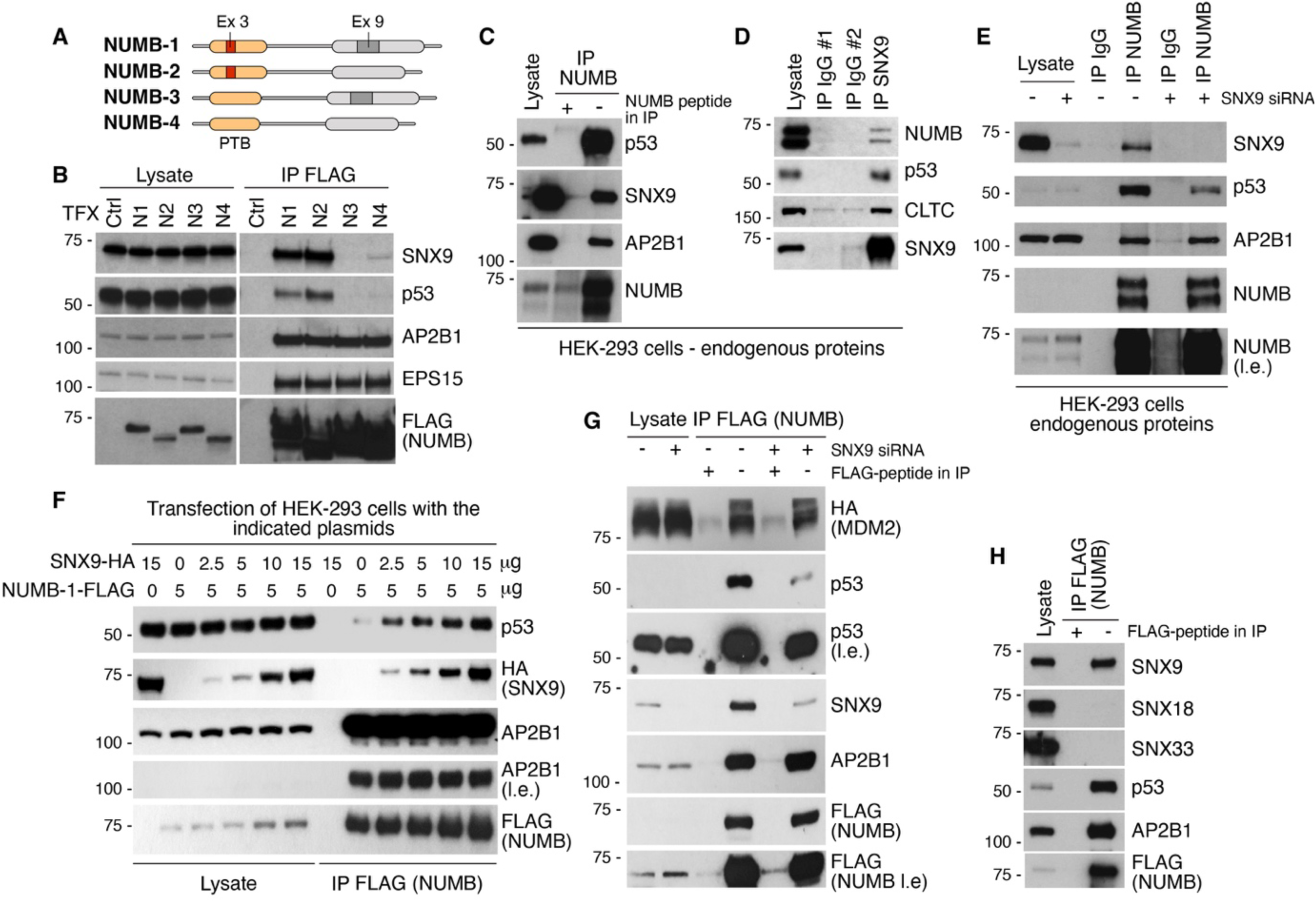
SNX9 specifically interacts with NUMB isoforms 1 and 2 and mediates the binding of NUMB to p53. **A.** Domain structure of NUMB isoforms. The positions of the two alternatively spliced exons (Ex3 and Ex9) and the PTB domain are shown. **B.** HEK-293 cells were transfected with vectors encoding the indicated FLAG-tagged NUMB isoforms (N1-N4) or empty vector (Ctrl). FLAG-NUMB was immunoprecipitated (IP) and immunoblotted (IB) as indicated on the right. Molecular weight markers are in kDa (here and all figures). **C.** Endogenous NUMB was IP from HEK-293 cells and IB as indicated. A peptide corresponding to the NUMB antibody epitope was used as negative control to compete antibody binding during the IP. **D.** HEK-293 cell lysates were IP using an anti-SNX9 antibody or irrelevant IgGs and IB as indicated. **E.** HEK-293 cells were silenced with SNX9 siRNA (+) or Ctrl siRNA (-), followed by IP with anti-NUMB or irrelevant IgGs and IB as indicated. **F.** HEK-293 cells were transfected with the indicated amounts of SNX9-HA or NUMB-1-FLAG and IB as shown (right). **G.** HEK-293 cells were silenced with SNX9 siRNA (+) or Ctrl siRNA (-) and transfected with NUMB-1-FLAG and MDM2-HA. Anti-FLAG IP was performed followed by IB as shown. **H.** HEK-293 cells transfected with NUMB-1-FLAG were IP using an anti-FLAG antibody and IB as shown. A FLAG peptide was used as negative control to compete antibody binding during the IP in **G** and **H**. In all panels, l.e., long exposure. Here and in subsequent figures, EPS15 and AP2B1 (AP2 complex subunit beta-1) are known NUMB (all isoforms) interactors, and CLTC (clathrin heavy chain 1) is a known SNX9 interactor.

From the evolutionary perspective, the emergence of Ex3 presents an intriguing puzzle. While NUMB itself is a metazoan innovation ^27, 38^, Ex3 was acquired later in the common ancestor of the chordate lineage, being present from the Cephalochordate and Tunicate subphyla onward and absent in invertebrates ^27^. This raises the question: why was a molecular function regulating p53 (Ex3) integrated into an endocytic protein like NUMB? We have previously shown that Ex3 is both necessary and sufficient to inhibit MDM2, as it retains this function even when engineered onto a completely unrelated scaffold ^1^. Was NUMB, then, merely an innocent bystander coopted to host a novel function, or does the “choice” of NUMB reflect a deeper mechanistic connection between endocytosis (and membrane traffic) and the regulation of p53 function?

In this study, we aimed to address this question by gaining a more detailed understanding of Ex3’s functions. Through proteomic analysis, we discovered that the Ex3-encoded sequence plays a central role in mediating multiple protein-protein interactions. A major interactor of the Ex3-encoded sequence is sorting nexin 9 (SNX9), a multi-functional scaffold protein involved in regulating endocytic processes and coordinating them with actin dynamics ^39^. SNX9 has also been implicated in cancer cell invasion and metastasis ^39^. The interaction between NUMB and SNX9 occurs exclusively at the plasma membrane (PM). Ex3 plays a multifaceted role in this interaction, as it interacts with phosphatidylinositol 4,5-bisphosphate (PI(4,5)P_2_) – thereby directing NUMB to the PM – and also with SNX9 in a bidentate interaction involving the canonical PTB binding groove of NUMB. At the PM, the SNX9:NUMB complex associates with p53, in a SNX9-dependent fashion, after which the complex is internalized, in a SNX9- and NUMB-dependent manner, routed to multivesicular bodies (MVB) and ultimately secreted in exosomes. The exosomal secretion of p53 exerts both cell-autonomous and non-cell-autonomous effects. In particular, exosomal p53 can be transferred to receiving cells, where it is translocated to the nucleus to initiate p53-dependent transcriptional and phenotypic changes. This, in turn, raises the intriguing possibility that exosome-mediated p53 secretion may contribute to the establishment of a tumor-suppressive microenvironment.

## 2. RESULTS

### 2.1 SNX9 functions as a molecular bridge between NUMB-1/-2 and p53

In mammals, NUMB exists in four major isoforms, generated by the alternative splicing of two exons, Ex3 and Ex9 (see Fig. 1A) ^37^. To gain insights into Ex3-associated functions, we performed a proteomic analysis to identify interactors that specifically bind to Ex3-containing NUMB isoforms (1 and 2), but not to Ex3-lacking isoforms (3 and 4) (Fig. 1A). HEK-293 cells were transfected with constructs encoding FLAG-tagged versions of the four isoforms. These tagged proteins were expressed at comparable levels and displayed the expected *in vivo* interactions with selected, known NUMB interactors (AP2B1 and EPS15, Fig. 1B). Cell lysates were then immunoprecipitated using an anti-FLAG antibody, and the resulting immunoprecipitates were analyzed by mass spectrometry (Fig. S1A). One of the most abundant specific interactors of NUMB-1/2 was SNX9, a protein with a known role in endocytic and trafficking processes (Fig. S1A and Fig. 1B). The interaction between NUMB and SNX9 was validated by co-immunoprecipitation (co-IP) of endogenous proteins in HEK-293 cells (Fig. 1C,D), as well as in MCF10A and HBL-100 cells (Fig. S1B,C).

Notably, endogenous p53 co-immunoprecipitated with NUMB-1/2 (Fig. 1B) as well as with endogenous SNX9 (Fig. 1D), suggesting that SNX9 could be part of a tripartite complex containing Ex3-containing NUMB isoforms and p53. NUMB silencing had no effect on the *in vivo* interaction between SNX9 and p53 (Fig. S1D), while SNX9 silencing significantly decreased the interaction between NUMB and p53 (Fig. 1E). In addition, SNX9 overexpression increased, in a dose-dependent fashion, the interaction between NUMB and p53 (Fig. 1F). The effect was specific, in that the *in vivo* interaction between NUMB and MDM2 – which is also known to interact specifically with the Ex3-encoded sequence of NUMB ^1^ – was not affected by SNX9 silencing (Fig. 1G, a rescue-control experiment is shown in Fig. S1E).

Finally, since the SNX9 subfamily of sorting nexins includes SNX18 and SNX33, we tested whether these two proteins also interact with NUMB. In transfected HEK-293 cells, no appreciable co-IP of SNX18 or SNX33 with FLAG-NUMB was detected (Fig. 1H).

These results suggest that Ex3-containing NUMB isoforms specifically interact *in vivo* with SNX9, which in turn binds to p53, acting as a bridge between NUMB and p53.

### 2.2 Molecular determinants mediating the NUMB:SNX9 interaction

To characterize the molecular interaction between NUMB and SNX9, we initially verified the involvement of the Ex3-encoded sequence *in vivo*. HEK-293 cells were transfected with constructs encoding FLAG-tagged full-length NUMB-1 or its PTB domain in its long (PTB-L) Ex3-containing or short (PTB-S) Ex3-lacking versions. By co-IP, we demonstrated that PTB-L was able to interact efficiently *in vivo* with SNX9 (and with p53), while PTB-S exhibited negligible interactions (Fig. 2A and Fig. S2A).

**Figure 2.**
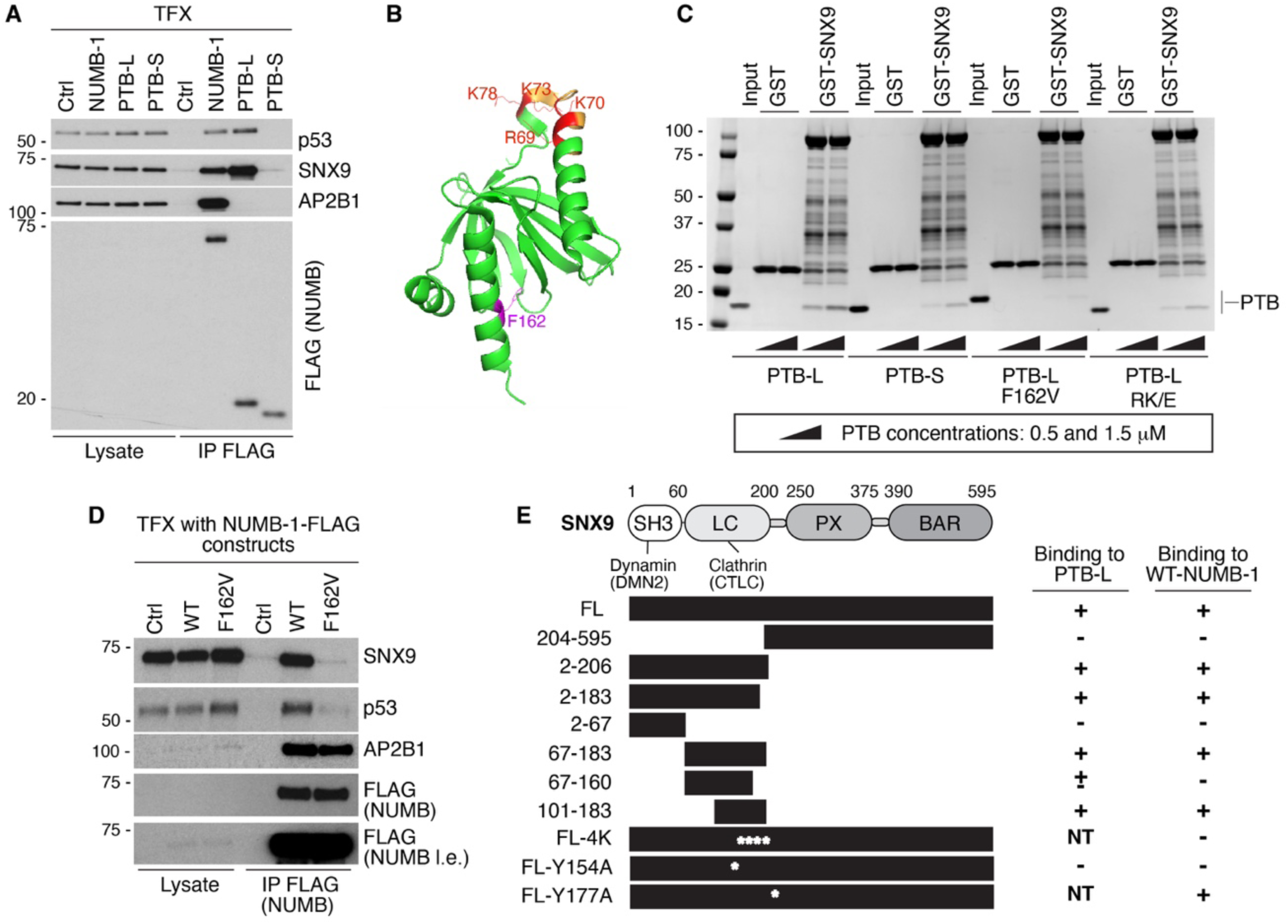
Characterization of the SNX9:NUMB interaction. **A.** HEK-293 cells were transfected (TFX) with the indicated NUMB/PTB FLAG-tagged constructs or empty vector (Ctrl), followed by IP with anti-FLAG antibody and IB as shown (right). **B.** Structure of PTB-L with the Ex3-encoded sequence indicated in yellow. The positions of key amino acids involved in protein-protein interactions are shown. In red, amino acids critical for MDM2 binding ^1^. In purple, the F162 residue in the PTB binding groove. **C.** *In vitro* binding assay with purified GST-SNX9 or GST alone as a negative control (top), immobilized on GSH beads, and purified NUMB-PTB proteins at increasing concentrations as indicated (bottom). After recovery, bound proteins were detected by Coomassie staining. **D.** HEK-293 cells were TFX with vectors encoding FLAG-tagged WT NUMB or the F162V mutant, or with EV (Ctrl). Anti-FLAG IP was performed followed by IB as shown (l.e., long exposure). **E.** Left, schematic of the SNX9 protein domain structure, with amino acid boundaries and binding sites of known interactors (dynamin and clathrin) indicated (top). The different SNX9 constructs tested are shown below (asterisks indicate positions of point mutations). Right, summary of the binding properties of SNX9 constructs to PTB-L and NUMB-1 (data are shown in Fig. S4 and S5).

PTB-L contains at least two known binding surfaces (Fig. 2B) – the Ex3-encoded sequence responsible for MDM2 binding (^1^ and Fig. S2B) and the canonical PTB-binding groove ^40^, also present in PTB-S. In *in vitro* binding assays with purified proteins, PTB-L exhibited dose-dependent binding to GST-SNX9 (Fig. 2C). Interestingly, PTB-S also bound to GST-SNX9, albeit less strongly than PTB-L. This result contrasts with the *in vivo* co-IP assays, which showed negligible binding between PTB-S and SNX9 (Fig. 2C, compare with Fig. 2A). Notably, mutagenesis of the two main binding sites in PTB-L (F162V in the PTB-binding groove and R69E/K70E/K73E/K78E – henceforth the RK/E mutant – in the Ex3-encoded sequence ^1^) revealed that F162 is critical for the interaction with SNX9 since the F162V mutant did not show detectable binding to purified SNX9 (Fig. 2C). Conversely, mutation of the Ex3-binding sites (RK/E mutant) decreased binding to SNX9 compared with PTB-L, but did not abrogate it (Fig. 2C). The strength of RK/E mutant binding to SNX9 was comparable to that of PTB-S, consistent with the lack of Ex3 in this version of the PTB (Fig. 2C). The critical role of the PTB-binding groove in SNX9 binding was confirmed by *in vivo* co-IP experiments (Fig. 2D) and size-exclusion chromatography (SEC) analysis of purified proteins (Fig. S3).

Finally, by performing co-IP, pull-down and *in vitro* binding assays with different purified SNX9 protein fragments or mutants, we mapped the optimal NUMB-PTB binding surface to the region spanning amino acids 101-183 (Fig. 2E and Fig. S4). This region is part of the low complexity (LC) domain of SNX9, which constitutes the least conserved region among the SNX9/18/33 family members (Fig. S5A,B).

Our data suggest that there are two SNX9 binding surfaces within the NUMB-PTB domain: a higher affinity surface within the canonical PTB-binding groove and a lower affinity surface within the Ex3-encoded sequence (see Fig. 2B). These binding surfaces should interact with distinct surfaces on SNX9, likely located within its LC domain. We hypothesized that an acidic region encompassing aa D162, D163, D166, and E167 (Fig. S5B) could be the binding surface for the Ex3-encoded sequence which is enriched in basic aa (Fig. 2B). This hypothesis is supported by the finding that a similar acidic aa-rich stretch in MDM2 mediates the binding with the Ex3-encoded sequence ^1^. In addition, given that the NUMB PTB-binding groove interacts with Y-containing peptides ^40^, we hypothesized that residue Y154 in the LC domain, which is unique to SNX9 and not present in the related SNX18 and SNX33 family members, might mediate this interaction (Fig. S5B). Residue Y177, which is present in both SNX9 and SNX18, could also be a candidate (Fig. S5B).

To test these hypotheses, we engineered SNX9 mutants in which the four acidic aa (D162, D163, D166, and E167) were mutagenized to K (henceforth, the SNX9-4K mutant) or Y154 and Y177 were individually mutagenized to A. These mutants were transfected into HEK-293 cells along with NUMB-1-FLAG, and their ability to interact was determined by co-IP. Both the SNX9-4K mutant and the SNX9-Y154A mutant were strongly impaired in their ability to co-IP with NUMB-1-FLAG, with the SNX9-Y154A mutant displaying negligible binding (Fig. S5C). Conversely, the SNX9-Y177A mutant was indistinguishable from WT SNX9 in its ability to co-IP with NUMB-1-FLAG (Fig. S5C). We confirmed that the impaired binding of the SNX9-Y154A mutant to NUMB was the result of a direct alteration of the interacting surface, as demonstrated by an *in vitro* binding assay (Fig. S5D). These results (summarized in Fig. 2E) support the hypothesis that the binding of SNX9 to NUMB-PTB is mediated by a bidentate interaction, in which both surfaces are required for optimal binding: one surface (NUMB-PTB-binding groove:SNX9-Y154) displays higher affinity than the other (NUMB-Ex3-polybasic stretch:SNX9-polyacidic stretch) (see also Fig. S5E).

### 2.3 Ex3-containing NUMB isoforms interact with SNX9 at the PM

Our data highlight a conundrum. The ability of NUMB to bind SNX9 *in vivo* appears to critically depend on the Ex3-encoded sequence, even though the canonical PTB-binding groove – present in all NUMB isoforms – represents the major binding surface for SNX9 (compare Fig. 2A, 2C and 2D). These findings suggest that the Ex3-encoded sequence determines the specificity for SNX9 interaction *in vivo* not simply via direct physical binding, but through indirect mechanisms.

To gain insights into Ex3-specific properties, we compared the subcellular localization of Ex3-containing (NUMB-1) and Ex3-lacking (NUMB-3) NUMB isoforms. Immunofluorescence (IF) imaging of FLAG-tagged NUMB-1 and NUMB-3 expressed in various cell types (MCF10A, Fig. 3A; HEK-293 and HBL-100, Fig. S6A,B), revealed clear differences in their subcellular distribution. NUMB-3 was detectable in numerous punctate structures distributed throughout the cytoplasm, most likely corresponding to endosomes ^41^. In contrast, NUMB-1 displayed a clear enrichment at the cell periphery, consistent with a predominantly PM localization. The NUMB-1-F162V PTB-binding groove mutant, which does not interact with SNX9 and p53 *in vivo* (see Fig. 2B,D), exhibited a similar localization to WT NUMB-1, at the cell periphery (Fig. 3A). In contrast, mutation of the Ex3-encoded sequence in the NUMB-1 RK/E mutant and NUMB-1 F71A/F72A/F75A/F76A mutant (F/A mutant) ^1^, resulted in a cytosolic punctate distribution similarly to the Ex3-lacking NUMB-3 isoform (Fig. 3A). These results suggest that the Ex3-encoded sequence determines a PM localization of NUMB. To confirm this finding, we explored the subcellular localization of endogenous NUMB isoforms. The selective KD of Ex3-containing isoforms (NUMB-1/2), yielded a residual NUMB staining, corresponding to isoforms 3-4, in punctate cytosolic structures (Fig. 3B). Conversely, the KD of isoforms 3-4 resulted in NUMB staining predominantly at the cell periphery consistent with a localization of isoforms 1-2 at the PM (Fig. 3B).

**Figure 3.**
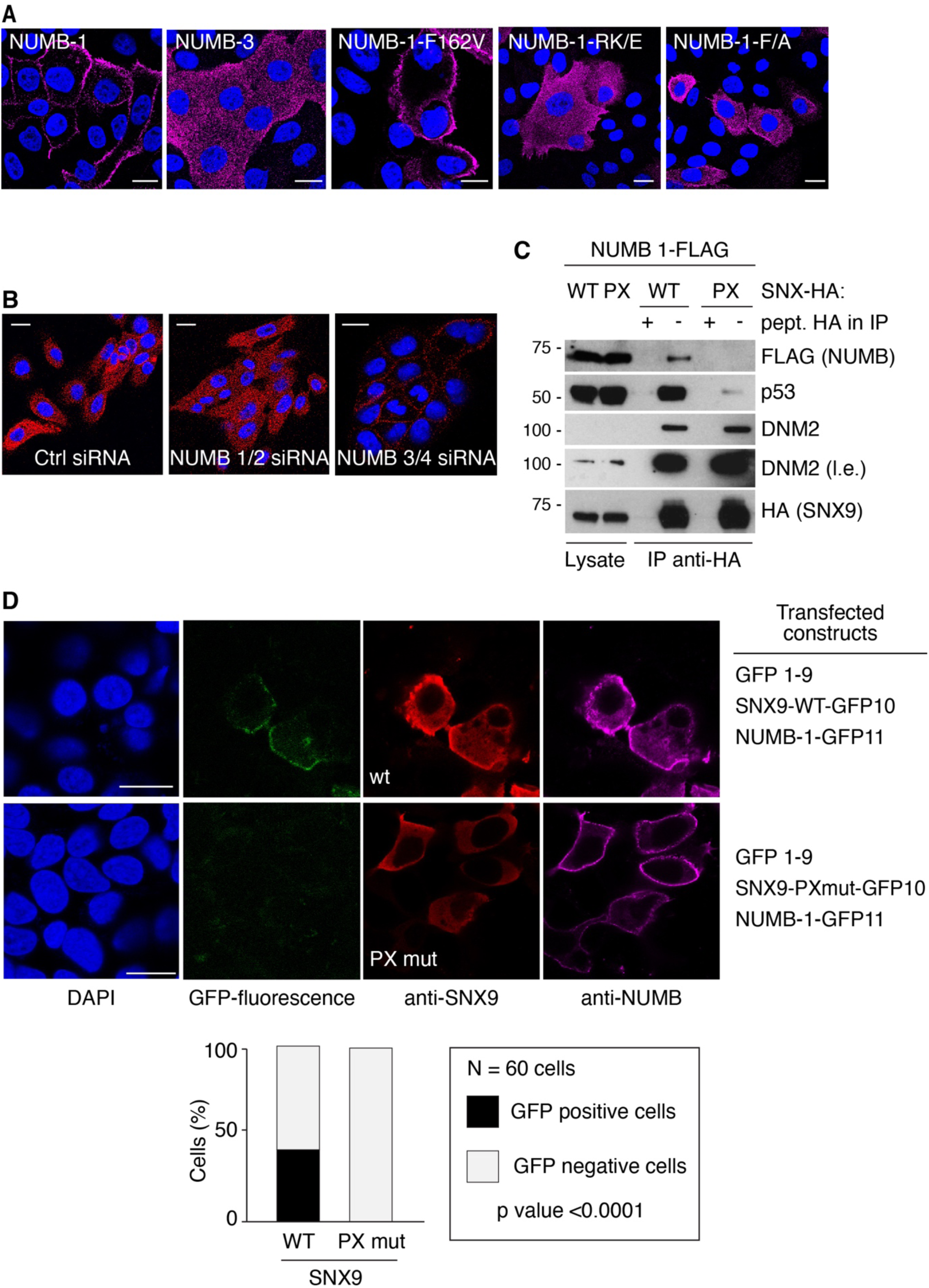
Ex3 localizes NUMB to the PM facilitating its interaction with SNX9. **A.** MCF10A cells were transfected with the indicated FLAG-tagged NUMB constructs. Transfected NUMB was visualized by immunofluorescence (IF) imaging with an anti-FLAG antibody (purple). Blue, DAPI counterstain. Bar, 20 μm. **B**. MCF10A cells were silenced with the indicated siRNAs and endogenous NUMB was visualized by IF imaging with anti-NUMB antibody (red). Blue, DAPI. Bar, 20 μm. **C.** HEK-293 cells were transfected with NUMB-1-FLAG and SNX9-HA WT or PXmut constructs (top). Anti-HA IP was performed followed by IB as indicated (right; l.e., long exposure). An HA peptide was used as negative control to compete with antibody binding during the IP. **D.** HEK-293 cells stably expressing the GFP 1-9 fragment and KO for SNX9 and NUMB, were transfected with the SNX9-WT/PXmut-GFP10 and NUMB-1-GFP11 constructs as indicated (right). Images of a representative IF analysis of SNX9 (red) and NUMB (purple) expression, GFP fluorescence and DAPI counterstain are shown. Bar, 20 μm. Quantification of the percentage of GFP-positive cells is shown below.

Based on these results, we hypothesized that the NUMB:SNX9 interaction occurs predominantly at the PM, explaining the specificity for Ex3-containing isoforms *in vivo*. To test this possibility, we engineered a SNX9 mutant (SNX9-PXmut) that fails to localize to the PM due to mutations (K366E/K367E) in its PX domain ^42^. As expected, this mutant retained its ability to bind NUMB-1 *in vitro* (Fig S6C,D), since its LC domain is unaltered. However, *in vivo,* its interaction with NUMB-1 (and with p53) was strongly impaired (Fig. 3C), supporting the hypothesis that the NUMB:SNX9 interaction occurs at the PM. To verify this finding, we used the tripartite-split GFP system to visualize the precise location of the NUMB:SNX9 interaction *in vivo* (Fig. S6E) ^43^. We observed reconstitution of the GFP-fluorescence signal at the PM only in the presence of NUMB-1 and SNX9-WT, while the combination of NUMB-1 with SNX9-PXmut did not produce any evident GFP-fluorescence signal (Fig. 3D). Therefore, the Ex3-encoded sequence of NUMB-1 appears to serve two main functions with respect to the NUMB:SNX9 interaction: i) it provides a low-affinity binding surface for SNX9, ii) it localizes NUMB-1 to the PM where it facilitates a “productive” interaction with SNX9. Since SNX9 localization at the PM is also required for the NUMB:SNX9 interaction (Fig. 3C,D), we propose a model in which both NUMB-1 and SNX9 bind independently to the PM where they can then interact with each other.

### 2.4 The NUMB:SNX9 complex associates with phospholipid bilayer structures and p53

Using liposomes as an *in vitro* model of phospholipid bilayer cellular membranes, we investigated how the Ex3-encoded sequence of NUMB-1 determines its interaction with the PM. We produced liposomes containing either PI(4)P or PI(4,5)P_2_ (hereafter referred to as PIP_2_) and tested their ability to bind recombinant PTB-L, PTB-S, and SNX9-WT proteins (Fig. S7A,B). Both PTB-L and SNX9-WT bound preferentially to PIP_2_-containing liposomes, whereas PTB-S exhibited no detectable binding to either type of liposome (Fig. 4A). Thus, the Ex3-encoded sequence of NUMB appears to mediate binding to both SNX9 and PIP_2_, a phosphoinositide enriched in the PM. This result might explain the preferential localization of NUMB isoforms 1 and 2 at the PM.

**Figure 4.**
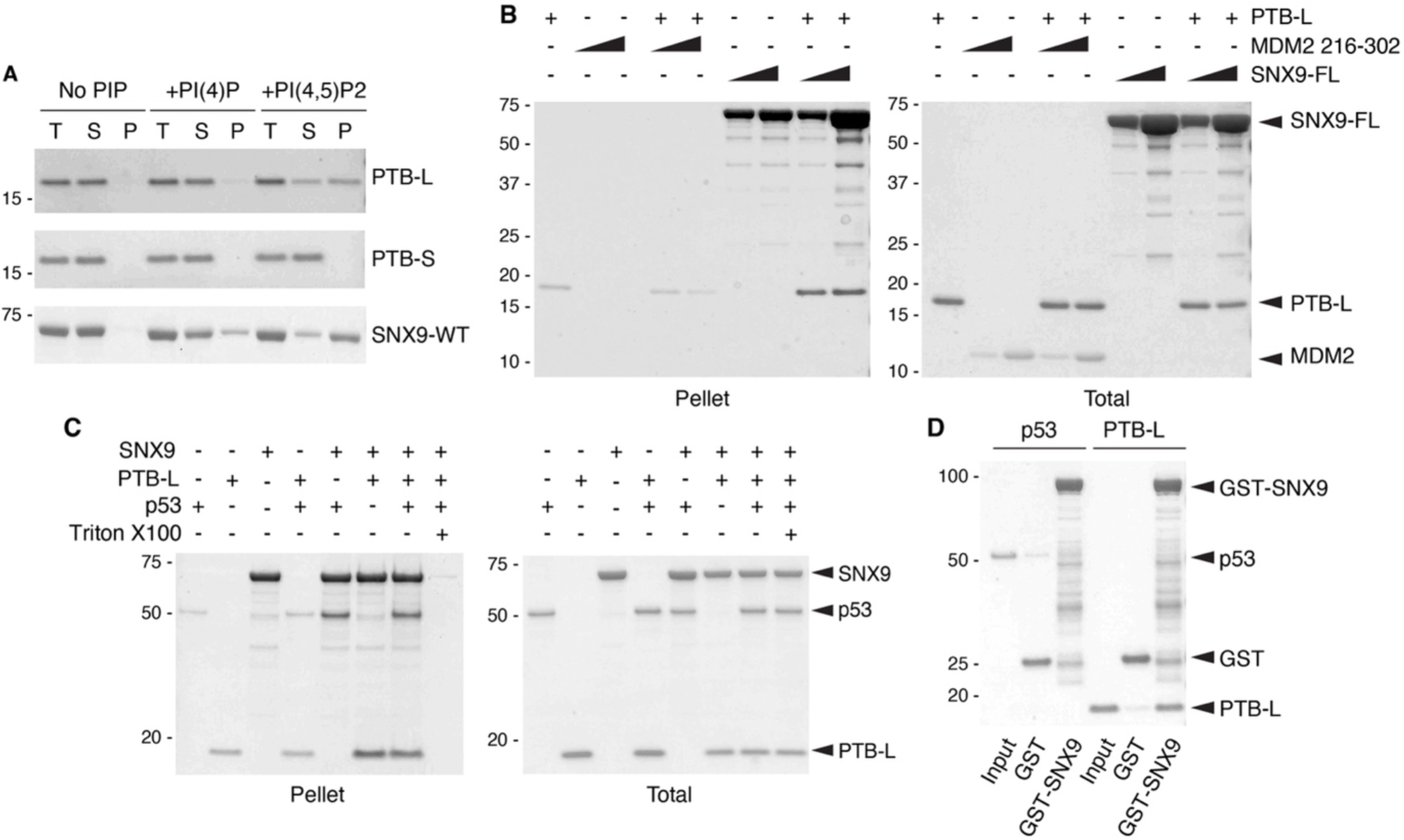
The NUMB:SNX9 complex associates with phospholipid bilayer structures and p53. **A.** Liposome binding assay with purified PTB-L, PTB-S, and SNX9-WT proteins (6 μM) and liposomes composed of PC/PS (No PIP) or PC/PS and either PI(4)P or PI(4,5)P2 as indicated. After centrifugation, the pellet (P) is enriched with liposomes and bound proteins. S: supernatant; T: total uncentrifuged sample. Bound proteins were detected by Coomassie staining. **B**. PIP_2_-liposome binding assay performed with purified PTB-L (10 μM), and increasing concentrations of SNX9 or MDM2^(216–302)^ (20 and 60 μM) as indicated. MDM2^(216-302)^ corresponds to aa 216-302 fragment harboring phosphomimetic mutations that increase binding to NUMB ^1^. Bound proteins were detected by Coomassie staining. The low staining intensity observed for MDM2 is attributable to the intrinsically low affinity of MDM2 for the Coomassie dye **C.** PIP_2_-liposome binding assay performed with purified SNX9 (10 μM), PTB-L (10 μM) and p53 (5 μM) as indicated. Triton-X100 was used to solubilize liposomes in negative controls. Bound proteins were detected by Coomassie staining. **D.** *In vitro* binding assay with purified GST-SNX9 or GST proteins immobilized on GSH beads and recombinant purified PTB-L (10 μM) and p53 WT (5 μM) proteins as indicated. Bound proteins were detected by Coomassie staining.

To further investigate this possibility, we treated NUMB-1-GFP-expressing MCF10A cells with ionomycin, which, by activating phospholipase C (PLC), depletes PIP_2_ from the PM ^44, 45^. This treatment resulted in the detachment of NUMB-1 from the PM, which, however, was prevented by cotreatment with the PLC inhibitor U73122 (Fig. S7C), supporting the idea that Ex3-containing NUMB isoforms localize to the PM by binding to PIP_2_.

Using the liposome binding assay, we also investigated how the interaction of PTB-L with biomembranes is influenced by its interaction with SNX9 or MDM2. The binding of both PTB-L and SNX9 to PIP_2_ liposomes increased when both proteins were present, whereas in the presence of MDM2, PTB-L binding to PIP_2_ liposomes was decreased (Fig. 4B, Fig. S7D). MDM2 alone did not show any detectable binding to liposomes. Thus, the presence of both PTB-L and SNX9 increases their avidity for PIP_2_, likely mediated through multiple interaction surfaces: PIP_2_:PTB-L, PIP_2_:SNX9, and SNX9:PTB-L. In contrast, the presence of MDM2 probably impairs the PTB-L:PIP_2_ interaction by masking the PIP_2_ binding surface within the Ex3-encoded sequence. Therefore, it is likely that the NUMB:SNX9 and NUMB:MDM2 complexes *in vivo* are mutually exclusive, with the former localized at the PM and the latter in the cytosol. This conclusion is supported by SEC data demonstrating that PTB-L in the presence of both MDM2 and SNX9 forms distinct dimeric complexes, PTB-L:MDM2 and PTB-L:SNX9, while a PTB-L:MDM2:SNX9 tripartite complex is not observed (Fig. S7E).

Next, we investigated the effects of the presence of biomembranes on the formation of the NUMB-PTB-L:SNX9:p53 tripartite complex. Purified recombinant p53 alone associated weakly with PIP_2_ liposomes, and the addition of PTB-L did not affect the binding of either p53 or PTB-L to liposomes, suggesting that these proteins interact with PIP_2_ independently of each other (Fig. 4C). In contrast, the addition of SNX9 substantially increased the individual binding of both p53 and PTB-L to PIP_2_ liposomes, whereas the presence of all three proteins together did not further increase their binding (Fig. 4C). Notably, in *in vitro* binding assays with purified recombinant GST-SNX9 and p53, no direct binding between SNX9 and p53 was detected, unlike with SNX9 and PTB-L (Fig. 4D). Since SNX9 can co-IP with p53 *in vivo* (Fig. 1D) and given the increased association of p53 with PIP_2_ liposomes in the presence of SNX9, we hypothesized that the SNX9:p53 interaction might require PIP_2_ binding. Specifically, the interaction of SNX9 with PIP_2_ might induce a conformational change in the protein, unmasking a p53 binding site. This possibility is compatible with the finding that SNX9-PXmut did not interact with p53 *in vivo* (Fig. 3C).

### 2.5 The NUMB:SNX9 complex localizes p53 at the PM

Our data suggest the existence of a NUMB-1/-2:SNX9:p53 complex at the PM. We tested this possibility by multiple imaging approaches. Initially, we adapted the IF protocol to use the mild detergent saponin, which permeabilizes the PM but not the nuclear membrane. This allows visualization of extranuclear p53, which would otherwise be obscured by the predominant nuclear p53 signal. Using this technique, we could readily identify areas of the PM in which the IF signals of the three proteins colocalized (Fig. S8A, see also Fig. S8B for specificity controls).

Next, we performed super-resolution fluorescence microscopy (direct stochastic optical reconstruction microscopy - dSTORM), staining the cells for SNX9 and p53 (Fig. 5A). A spatial analysis of the signals revealed a higher percentage of co-localization in areas close to the PM compared with the rest of the cytoplasm (Fig. 5A).

**Figure 5.**
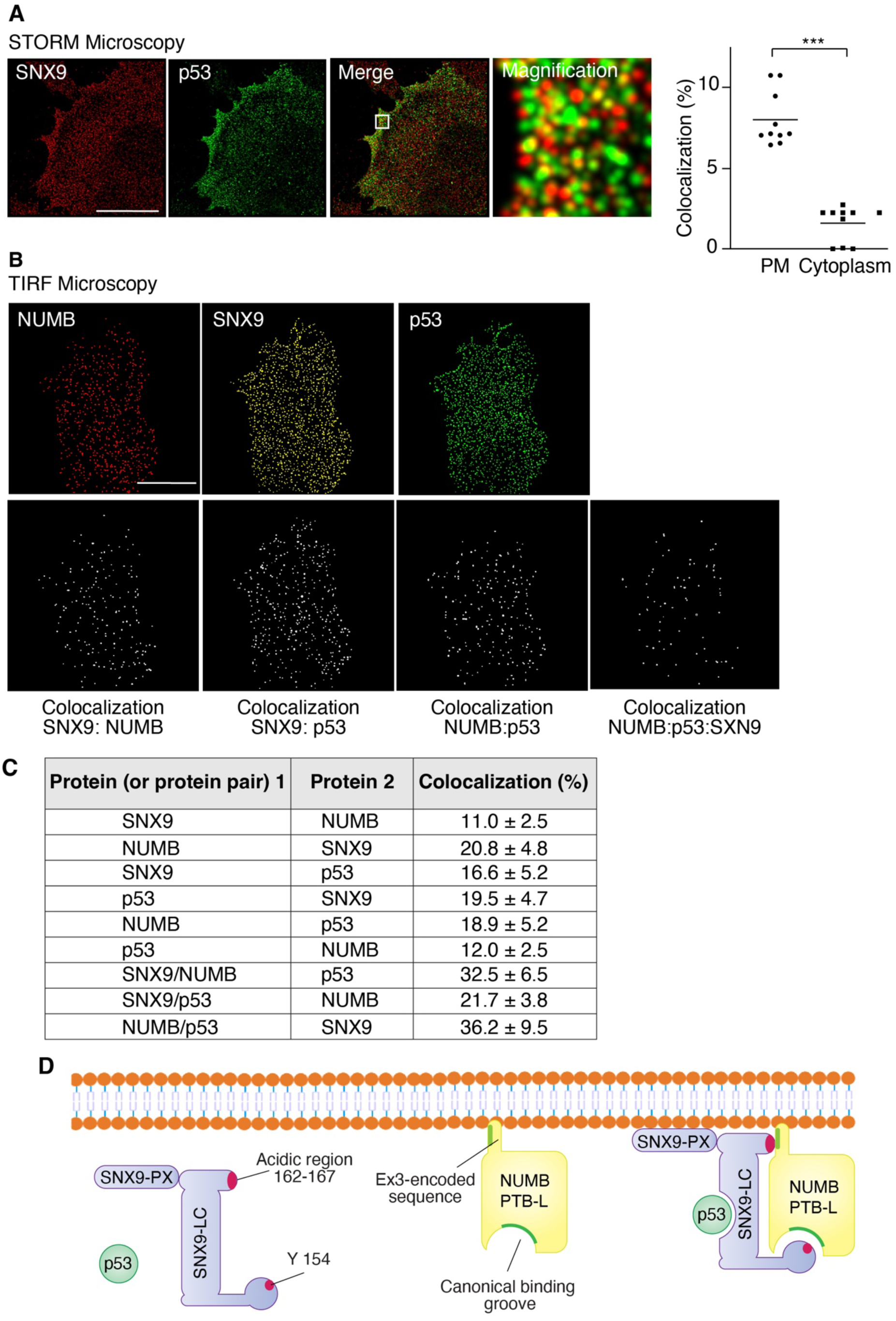
The interaction between Ex3-containing NUMB isoforms, SNX9 and p53 occurs at the PM. **A.** Dual color dSTORM images with anti-SNX9 (red) or anti-p53 (green) antibodies in HBL-100 cells. Bars, 20 μm. The percentage colocalization of the two channels in the vicinity of the PM (defined as the region within 2 μm from the cell edge) and in the cytoplasm is shown on the right as the mean ± SD (N = 10 cells). **B.** A.M.I.CO image analysis of NUMB, SNX9 and p53 spot distribution at the basal membrane of representative TIRF images of HBL-100 cells (top panels). Colocalization is represented by white dots in the bottom panels. Bars, 20 μm. **C**. Summary of the percentage colocalization of the indicated proteins in the TIRF experiments shown in **B**. In each row the % of colocalization of the indicated protein 1 (or protein pair 1) with protein 2 is shown. Results are expressed as the mean ± SD (N = 10 cells). **D.** Cartoon depicting the NUMB:SNX9:p53 tripartite complex assembled at the PM (only the relevant portions of the proteins are shown). Binding to the PM is mediated through the SNX9-PX domain and the Ex3-encoded sequence of NUMB-PTB-L. SNX9 and PTB-L interact through a bidentate interaction involving two surfaces in the low-complexity (LC) domain of SNX9, and the Ex3-coding sequence and the canonical PTB-binding groove on NUMB (lower and higher affinity surfaces, respectively). Upon binding to the PM, conformational changes in the SNX9 LC domain allow it to bind p53.

Finally, by total internal reflection fluorescence (TIRF) microscopy, we investigated the co-localization of p53, SNX9 and NUMB at or near the cell surface (Fig. 5B). In pairwise comparisons, 10 to 20% of the PM membrane pools of the individual proteins associated with each other (Fig. 5C). In three-way comparisons, 20-35% of each protein localized to dots containing the other two proteins (Fig. 5C).

The sum of our data supports the existence of a NUMB:SNX9:p53 tripartite complex localized at the PM, mediated through the interactions depicted in the model in Fig. 5D.

### 2.6 p53 is packaged into exosomes in a NUMB and SNX9-dependent manner

The observation that a pool of WT endogenous p53 is localized at the PM suggested that it could have non-canonical functional roles at this location. Our investigations uncovered the presence of SNX9, NUMB and p53 in extracellular vesicles (EVs) isolated from HEK-293 cells (Fig. 6A, see Fig. S9A-C for quality controls). Notably, co-IP experiments revealed that SNX9 binding to both p53 and NUMB was maintained in these EVs (Fig. 6B).

**Figure 6.**
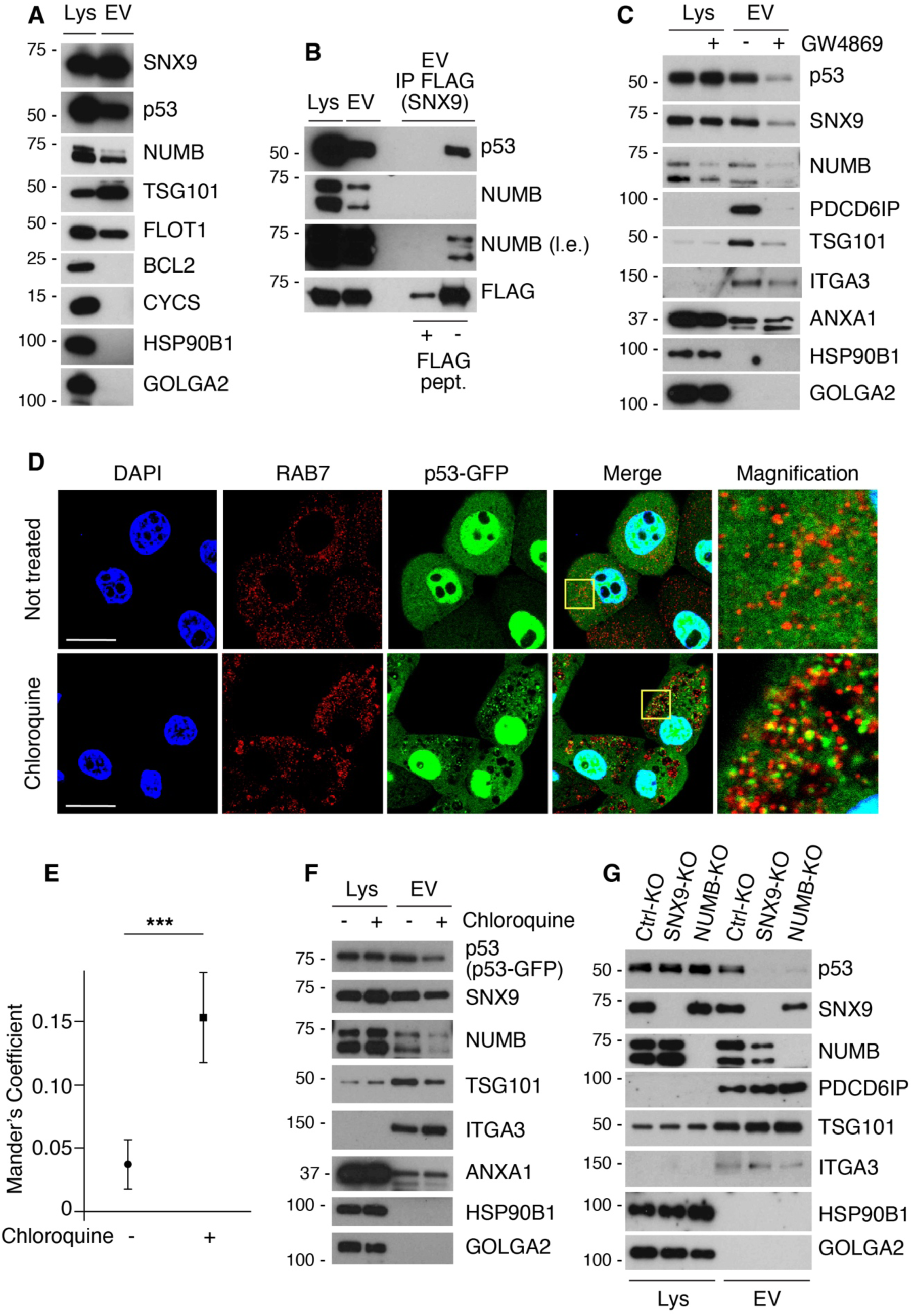
NUMB and SNX9 facilitate p53 packaging into exosomes. **A.** Extracellular vesicles (EV) were purified from the conditioned medium of HEK-293 cells and IB as shown (right). Lys, total cellular lysate (here and in all relevant panels). The quality of the EV purification was assessed using positive EV markers TSG101 and Flotilin-1 (FLOT1) and negative EV markers (BCL2; cytochrome c, CYCS; HSP90B1 also known as GRP94; GOLGA2 also known as GM130). **B.** HEK-293 cells were transfected with SNX9-FLAG and EVs were purified from conditioned medium. Anti-FLAG IP of isolated EVs was performed, followed by IB with the indicated antibodies (right; l.e., long exposure). A FLAG peptide was used as negative control to compete antibody binding during the IP. **C.** HEK-293 cells were treated with GW4869 (20 μM for 24 h). EVs were purified from conditioned medium and analyzed by IB as indicated (right). PDCD6IP (Alix) and TSG101 are enriched in exosomes. ANXA1 (annexin 1) and ITGA3 (integrin α3) are enriched in microvesicles. **D.** MCF10A cells stably expressing p53-GFP (green) were treated with chloroquine (50 μM for 24 h) or vehicle control. Anti-RAB7 (red) IF analysis was performed to visualize late endosomes. Blue. DAPI counterstain. Merged and magnified images are shown on the right. Bar, 20 μm. **E.** Quantification of RAB7 and p53-GFP colocalization, using Mander’s coefficient, in the experiment shown in panel D. A total of 8 fields from untreated cells and 12 fields from chloroquine-treated cells were analyzed by confocal microscopy. **F.** EVs were purified from the conditioned medium of the cells in panel D and IB as indicated. **G.** EVs were purified from the conditioned medium of Ctrl-KO, SNX9-KO or NUMB-KO HEK-293 cells as indicated on top and IB as shown (right).

EVs are produced by two major pathways: (i) budding of microvesicles directly from the PM, and (ii) secretion of exosomes, which are generated via the endosomal pathway and released upon fusion of multivesicular bodies with the PM. To distinguish between these two pathways, we treated cells with GW4869, a widely used inhibitor of exosome biogenesis ^46, 47^. This treatment led to a marked reduction in the levels of p53, NUMB and SNX9 in EVs, as well as PDCD6IP (Alix) and TSG101 – well-established exosome markers. In contrast, the levels of ANXA1 (annexin 1) and ITGA3 (integrin α3), which are preferentially incorporated into microvesicles ^48, 49^, were only marginally affected (Fig. 6C). Furthermore, in MCF10A cells stably expressing p53-GFP, we observed an accumulation of p53-GFP in cytoplasmic puncta following treatment with chloroquine: a lysosomotropic agent that inhibits endosomal acidification and disrupts exosome biogenesis via the endosomal pathway (Fig. 6D,E). Blockade of the endosomal pathway was confirmed by the co-localization of p53-GFP puncta with RAB7, a marker of late endosomes (Fig. 6D,E). Chloroquine treatment also decreased the levels of p53-GFP in purified EVs, along with NUMB, SNX9 and the exosomal marker TSG101, while having no effect on the microvesicle markers ITGA3 and ANXA1 (Fig. 6F). These findings indicate that p53 is secreted from cells via exosomes.

Mechanistically, the packaging of p53 into exosomes was dependent on both SNX9 and NUMB, since EVs purified from SNX9-KO or NUMB-KO cells exhibited substantially reduced levels of p53, while other exosome and microvesicle markers were not affected (Fig. 6G, see also Fig. S9D for additional controls). Finally, we quantified the amount of cellular p53 secreted via exosomes in MCF10A cells. ∼10% of the steady-state p53 content is packaged into exosomes and secreted per hour (Fig. S9E). Additionally, we estimated that MCF10A cells produce approximately 60 EVs per cell per hour. Based on these values, we calculated that each EV contains a minimum ∼100 p53 molecules (Fig. S9E).

### 2.7 Biological effects of p53 secretion via exosomes

The exosomal secretion of p53 has the potential to influence both the donor and recipient cells. In donor cells, the export of p53 would be expected to decrease its intracellular levels and downregulate its activity in a cell-autonomous manner. Conversely, in recipient cells, exosome-mediated delivery of p53 could increase its levels and activity in a non-cell-autonomous manner.

To investigate these possibilities, we used MCF10A cells as a more physiological, non-tumorigenic mammary epithelial cell model. The effects of p53 export on donor cells (cell-autonomous effects) were examined by inhibiting its exosomal secretion through SNX9 KO. We first confirmed that p53 levels in EVs derived from MCF10A-SNX9-KO cells were reduced compared with control cells (Fig. S10A). Notably, basal intracellular levels of p53 were increased following SNX9-KO, and this effect became more pronounced after treatment with the genotoxic drug etoposide (Fig. 7A). In response to etoposide-induced DNA damage, p53 is known to be phosphorylated (e.g., at Ser15) and stabilized, leading to increased protein levels and transcription of its downstream target genes ^50^. Consistent with this, we observed that elevated p53 levels in MCF10A-SNX9-KO cells were accompanied by increased transcription of the p53 target genes MDM2 and CDKN1A (encoding p21^Cip1^), particularly in etoposide-treated cells (Fig. 7A,B). Thus, SNX9-mediated exosomal secretion of p53 appears to downregulate p53 levels and activity in the donor cell.

**Figure 7.**
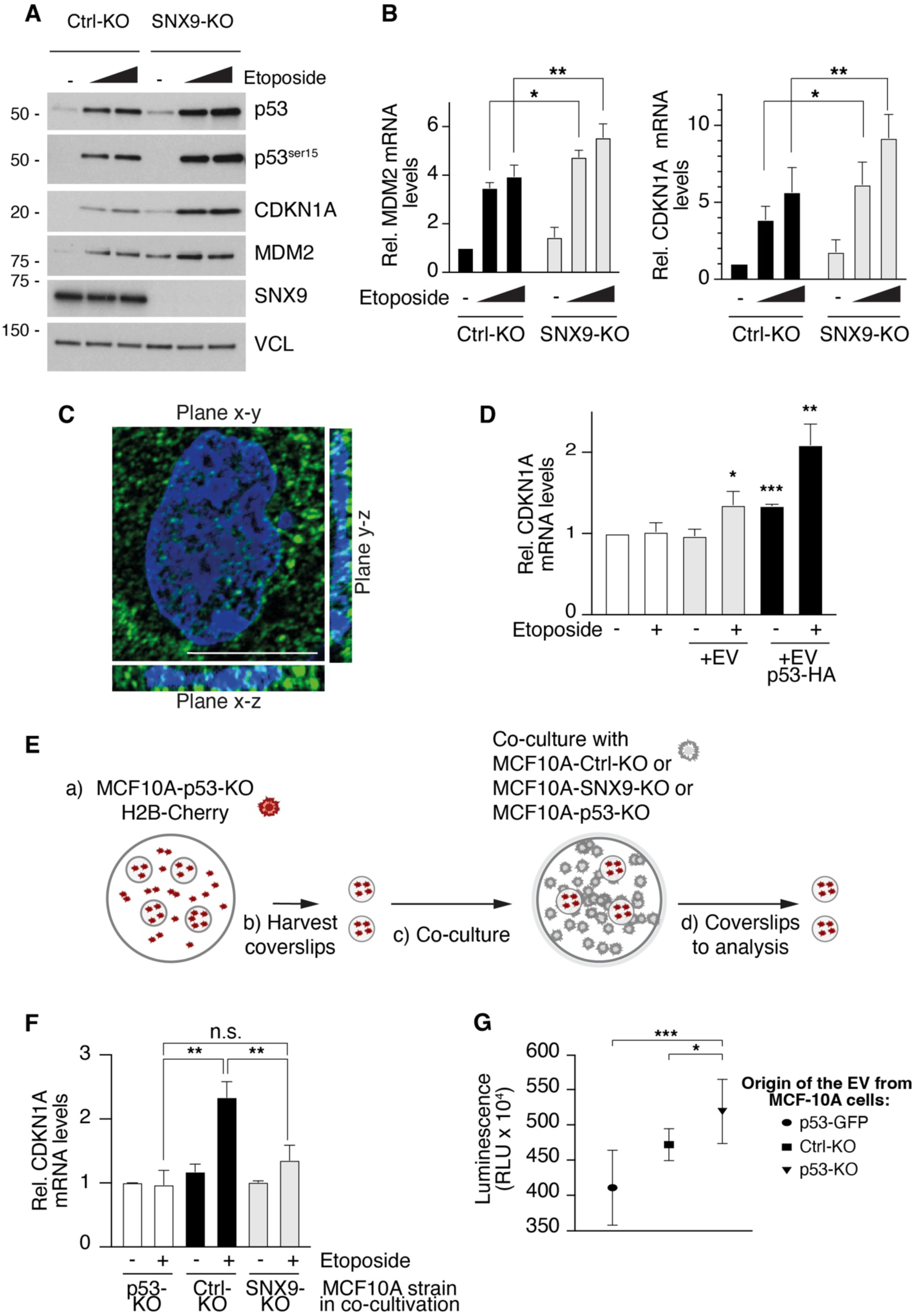
Cell-autonomous and non-cell-autonomous effects of p53 secreted via exosomes. **A.** Ctrl-KO and SNX9-KO MCF10A cells were treated with etoposide at 25 and 50 μM (indicated by triangles) and analyzed by IB for total and phosphorylated (p53^ser15^) p53, and the p53 target genes MDM2 and CDKN1A. Vinculin (VCL), loading control. **B**. RT-qPCR analysis of MDM2 and CDKN1A expression in the samples described in “A”. Data are from three independent experiments and expressed as mean ± SD. *, p<0.05; **, p<0.01. **C.** 3D reconstruction of MCF10A-p53-KO recipient cells treated for 8 h with EVs purified from the conditioned medium of HEK-293 cells transfected with p53-HA. Recipient cells were treated with etoposide (50 μM for 8 h) and analyzed by IF using an anti-p53 antibody (green) and DAPI (blue). Bar, 20 μm. **D.** MCF10A-p53-KO recipient cells were treated for 8 h with etoposide (50 μM) and EVs purified from HEK-293 WT (EV) or HEK-293 p53-HA (EV p53-HA) cells as indicated. After treatment cells were harvested and analyzed by RT-qPCR for CDKN1A levels. Data are from three independent experiments and expressed as mean ± SD. * p<0.05, ** p<0.01 and *** p<0.001 *vs*. same condition in cells not treated with EVs (only the most relevant statistical comparisons are shown). **E.** Scheme of the co-culture experiment shown in in panel F. a) MCF10A-p53-KO-H2B-Cherry cells were plated onto coverslips. b) After 24 h, coverslips were harvested and seeded (c) in plates in which the indicated cell lines had been previously seeded. Cells were then treated with etoposide (50 μM) or mock-treatment for 8 h. d) Coverslips were harvested and analyzed for purity of H2B-Cherry labeled cells (Fig. S10D) and for the levels of CDKN1A mRNA (panel F). Details are in Materials and Methods. **F**. RT-qPCR analysis of CDKN1A levels in harvested H2B-Cherry labeled MCF10A-p53-KO cells, co-cultured as described in panel “**E**”. Data are from three independent experiments and expressed as mean ± SD. **, p<0.01; n.s., not significant (only the most relevant statistical comparisons are shown). **G**. SAOS2 cells were treated with EVs purified from the indicated MCF10A cell lines (see experimental scheme in Fig. S10F), and cell viability/growth was assessed indirectly by quantifying intracellular ATP levels using a luminescence-based assay. Data are from ten samples/condition from two independent experiments and expressed as mean ± SD. One-way ANOVA test: *, and ***, p < 0.05 and < 0.001, respectively, *vs*. SAOS2 treated with EVs derived from MCF10A-p53-KO cells.

To investigate the effects of exosome-mediated p53 transfer on recipient cells, we used MCF10A-p53-KO cells (Fig. S10B) to specifically uncover non-cell-autonomous p53 functions. We verified that these cells do not exhibit p53 transcriptional activity, as evidenced by the negligible CDKN1A protein and transcripts levels, even after etoposide treatment (Fig. S10B). MCF10A-p53-KO cells were then treated with EVs purified from HEK-293 donor cells transfected individually with tagged SNX9-FLAG, NUMB-1-GFP or p53-HA constructs. All three proteins, transferred via exosomes, were clearly detectable in recipient cells (Fig. S10C). Notably, a fraction of secreted p53-HA appeared to be imported into the nuclei of recipient cells (Fig. 7C) and was transcriptionally active, as evidenced by increased CDKN1A transcript levels in MCF10A-p53-KO recipient cells treated with EVs containing p53-HA, in both basal and etoposide treated conditions (Fig. 7D). Interestingly, increased CDKN1A transcript levels were also observed in recipient cells treated with EVs from WT donor cells expressing physiological p53 levels, albeit to a lesser extent (Fig. 7D).

Next, we investigated the exosomal transfer of p53 directly between cells in a co-culture experiment. H2B-Cherry labeled MCF10A-p53-KO cells were used as recipient cells, co-cultured with an excess of MCF10A-Ctrl-KO, MCF10A-SNX9-KO or MCF10A-p53-KO cells used as donor cells (see scheme in Fig. 7E and controls in Fig. S10D). As expected, co-culture with MCF10A-p53-KO donor cells was unable to rescue the p53 response to etoposide in H2B-Cherry labeled MCF10A-p53-KO recipient cells (Fig. 7F). Only when MCF10A-p53-KO recipient cells were co-cultured with MCF10A-Ctrl-KO donor cells did we observe a significant increase in CDKN1A transcript levels following etoposide treatment, indicative of the transfer of functional p53 from MCF10A-Ctrl-KO to H2B-Cherry-MCF10A-p53-KO cells (Fig. 7F). In contrast, co-culture with MCF10A-SNX9-KO donor cells resulted in a significantly impaired p53 response in recipient cells (Fig. 7F), in agreement with the reduced exosomal export of p53 from MCF10A-SNX9-KO cells (Fig. S10A).

Finally, we investigated the phenotypic effects of p53 transfer via EVs. MCF10A cells were found to be a suboptimal model for studying the impact of altered p53 levels, as we observed high tolerance to both p53 KO and overexpression. Therefore, we selected SAOS2 cells, a p53-null osteosarcoma cell line known to undergo cell cycle arrest and/or apoptosis upon p53 restoration ^51, 52^. First, we confirmed that treatment with EVs derived from MCF10A-p53-GFP cells resulted in a detectable p53 signal, both in the cytoplasm and the nucleus of recipient SAOS2 cells (Fig. S10E). We then treated SAOS2 cells with EVs derived from MCF10A-Ctrl-KO, MCF10A-p53-KO or MCF10A-p53-GFP cells, and monitored cell growth over four days (see scheme in Fig. S10F). SAOS2 cells treated with EVs derived from MCF10A-Ctrl-KO cells – and more prominently with EVs from MCF10A-p53-GFP cells – exhibited a significant reduction in growth compared to those treated with EVs from MCF10A-p53-KO cells (Fig. 7G). These findings indicate that p53 transferred via exosomes can influence the behavior of recipient cells in a dose-dependent manner.

Together, these data indicate that exosomal secretion of WT p53 confers non-cell-autonomous p53 activity to recipient cells.

## 3. DISCUSSION

Here, we uncover a novel function for Ex3-containing NUMB isoforms in modulating the p53 regulatory network, beyond their established function in stabilizing p53 by inhibiting MDM2 ^1, 2, 27^. This regulatory mechanism is closely intertwined with the well-characterized roles of NUMB and SNX9 as endocytic adaptors governing trafficking. Our data demonstrate that SNX9 is essential for retaining p53 at the PM and that the assembly of the NUMB:SNX9:p53 tripartite complex is necessary for its internalization, leading to its incorporation into exosomes.

NUMB is therefore implicated in two distinct complexes that regulate p53 with opposing effects: through its interaction with MDM2, NUMB stabilizes p53 and increases its intracellular levels, whereas through its association with SNX9 at the PM, it promotes exosomal secretion of p53, thereby reducing intracellular p53 levels. Notably, our data reveal that the NUMB:SNX9 and NUMB:MDM2 complexes are mutually exclusive, with their relative stoichiometry likely determined by competition between the PM and MDM2 for binding to the Ex3-encoded sequence. The precise mechanism governing the dynamic balance between these two complexes – and the selective binding of Ex3-containing NUMB isoforms to either MDM2 or the PM – remains unresolved. However, under steady-state conditions, it is plausible that the availability of Ex3-binding sites at the PM (likely mediated by PIP_2_) vastly exceeds the availability of MDM2 in the cytosol, as confirmed by the predominant localization of NUMB-1/2 at the PM. Regulatory events, potentially involving phosphorylation, may disrupt the NUMB:PM interaction, thereby enabling NUMB to engage with MDM2. Indeed, phosphorylation-induced detachment of NUMB from the PM is a conserved feature throughout evolution ^37, 53^.

The NUMB:SNX9-mediated exosomal secretion of p53 is of considerable magnitude, accounting for approximately 10% of the intracellular p53 pool per hour. This release has significant implications for both cell-autonomous and non-cell-autonomous p53 functions. In the donor cell, exosome-mediated loss of p53 contributes to maintaining low levels of the protein, a process previously attributed primarily to the futile p53 cycle driven by the negative feedback loop between p53 and MDM2 ^54, 55^. However, exosome-mediated delivery of p53 also modulates p53-dependent responses in recipient cells, where exogenous p53 can be transported to the nucleus to activate p53-dependent transcriptional responses. To our knowledge, this represents the first demonstration that WT p53 can exert a non-cell-autonomous function through direct intercellular transfer.

We propose that this novel role of p53 can be best understood in the framework of the cell competition theory. Cell competition is a conserved fitness-sensing process whereby more robust cells eliminate less fit neighboring cells, a phenomenon occurring across various developing and adult tissues ^56–58^. Furthermore, cell competition can act as both a tumor-suppressive and tumor-promoting force, playing a pivotal role in cancer initiation and progression ^56^. p53 is emerging as a key player in cell competition ^59–63^, projecting a major role of this function of p53 in cancer. In this context, it is noteworthy that p53 haploinsufficiency might be sufficient to generate “winner” cell clones in a cell competition setting, as demonstrated by the increased cancer susceptibility of p53+/− mice, despite retention of a WT p53 allele in over half of the tumors ^64, 65^. In contrast, “super p53” mice, which harbor additional copies of p53, exhibit increased tumor resistance compared to WT animals ^66^. The loss of a single p53 allele can also trigger chronic inflammatory, DNA damage, and oxidative stress responses that promote carcinogenesis ^67^. This situation could be particularly relevant in the context of pro-tumorigenic tumor-stroma interactions. For instance, in a mouse model of prostate carcinogenesis, the loss of a single p53 allele in stromal fibroblasts led to their proliferation and accelerated tumor growth, even though the tumor epithelium retained WT p53 ^68^.

Based on our findings, we propose that exosomal secretion of WT p53 may represent a protective mechanism to preserve a tumor-suppressive environment. This process could compensate for the loss of heterozygosity of this key tumor suppressor gene in individual cells, which might otherwise gain a proliferative advantage and initiate tumorigenesis. Of note, exosomes carrying mutant p53 have been shown to promote tumor progression and induce the transformation of fibroblasts into a cancer-associated phenotype ^69^. According to our hypothesis, this would not represent a neomorphic function of mutated p53, but the hijacking of a physiological mechanism of tumor suppression.

## 4. EXPERIMENTAL SECTION

### Cells and chemicals

The MCF10A, HEK-293, SAOS2 and HBL-100 cell lines were cultured at 37°C in a humidified atmosphere with 5% CO_2_, using specific culture media: MCF10A in a 1:1 mixture of DMEM (Euroclone) and Ham’s F12 (Thermo Fisher Scientific), supplemented with 5% horse serum (Euroclone), 2 mM L-glutamine (Euroclone), 10 µg/mL human insulin (Euroclone), 0.5 µg/mL hydrocortisone (Euroclone), 100 ng/mL cholera toxin (Sigma-Aldrich), 50-100 µg/mL penicillin/streptomycin (Euroclone) and 20 ng/mL hEGF (CiteAb); HEK-293 in DMEM with 10% South American fetal bovine serum (FBS) (Euroclone), 2 mM L-glutamine and 50-100 µg/mL penicillin/streptomycin; SAOS2 in McCoy’s 5A (Sigma-Aldrich), with 15% South American FBS, 50-100 µg/mL penicillin/streptomycin, and 2 mM L-glutamine; HBL-100 in McCoy’s 5A, supplemented with 10% North American FBS (Euroclone), 50-100 µg/mL penicillin/streptomycin, and 2 mM L-glutamine. Expi293 cells (Gibco) were cultured in Expi293 Expression Medium (Gibco) according to the manufacturer’s instructions, in suspension and under agitation at 37 °C in a humidified atmosphere with 8% CO₂. All human cell lines were authenticated at each batch freezing by STR profiling (StemElite ID System, Promega). Mycoplasma tests were routinely performed on all cell lines. Transfection of HEK-293 cells was performed with the calcium phosphate method or with polyethylenimine (PEI) (Polysciences) using a ratio of 2.5 µg PEI per 1 µg DNA. The solution was incubated at RT for 10 min and then added to cells, followed by incubation for 6 h or overnight. MCF10A and HBL-100 cells were transfected with the JetOptimus Kit (Polyplus Transfection) according to the manufacturer’s instructions. Cells were harvested after 24-48 h. Expi293 cells were transfected with the Expifectamine 293 transfection kit (Gibco) according to the manufacturer’s instructions. HEK-293 GFP1-9 cells, shown in Fig. 3D, were generated by transfecting HEK-293 cells with the pcDNA-GFP1-9 plasmid (see section “Engineering of constructs”). Cells were selected with G418 (AdipoGen) (300 µg/ml), and clonal populations isolated by limiting dilutions were subsequently transfected for CRISPR-Cas9-mediated knockout (KO) of SNX9 and NUMB. A bulk population was obtained by puromycin (AdipoGen) selection (1 μg/ml). MCF10A p53-GFP cells were generated by lentiviral infection with the PLVX-GFP-p53 construct, followed by puromycin selection (1 μg/ml). Individual clones were isolated by limiting dilutions. The integrity of the p53-GFP fusion protein was confirmed by WB and IF, based on the co-localization of the GFP signal with p53 staining. MCF10A-p53-KO-H2B-Cherry cells were generated by retroviral infection of MCF10A-p53-KO cells with the plasmid H2B-PamCherry. Cells were selected with puromycin (1 μg/ml) and then sorted using a flow cytometer to isolate the Cherry-positive population.

Chemicals were: FLAG peptide, cat. F3290 (Merck Life Science); HA peptide, cat. 11666975001 (Merck Life Science); NUMB peptide corresponding to amino acids 537-551 of hNUMB (Genscript); Etoposide, cat. E1383 (Merck Life Science); Trehalose Dihydrate, cat. T9531 (Merck Life Science); Chloroquine, cat. C6628 (Merck Life Science); Ionomycin, cat. I0634 (Merck Life Science); Proteinase K, P4850 (Merck Life Science); MG132, cat. 474790 (Merck Life Science); U73122, cat. 6756 (Merck Life Science); GW4869, cat. S7609 (Selleck Chemicals); brain PI(4,5)P₂, cat. 840046P; brain PI(4)P, cat. 840045P; brain PS, cat. 840032C; brain PC, cat. 840053C (all from Merck Life Science).

### Antibodies

Primary antibodies (Ab) for immunofluorescence (IF) were directed against: NUMB (AB21, a mouse monoclonal Ab against amino acids 537–551 of hNUMB ^2^); SNX9 (15721-1-AP, Proteintech); FLAG (2368, Cell Signaling); p53 (DO-1, Santa Cruz Biotechnology for all the IF experiments, except for Fig S8A in which the AF1355, R&D Systems, was used); RAB7 (D95F2, Cell Signaling Technology). Fluorochrome-conjugated secondary Ab were from Jackson ImmunoResearch Laboratories.

Ab for immunoblot (IB) were directed against: NUMB (AB21, ^2^ and C29G11, Cell Signaling Technology); SNX9 (15721-1-AP, Proteintech); FLAG (2368, Cell Signaling Technology); p53 (DO-1, Santa Cruz Biotechnology and AF1355, R&D Systems); VCL (mouse monoclonal, Sigma-Aldrich); MDM2 (OP46, Calbiochem); AP2B1 (PA11066, Thermo Fisher Scientific); EPS15 (C-20, Santa Cruz); CLTC (610499, BD Biosciences); HA (C29F4, Cell Signaling Technology); SNX18 (21946-1-AP Proteintech); SNX33 (A305755AT, Thermo Fisher Scientific); DNM2 (AB3457, Abcam); TSG101 (AB30871, Abcam); FLOT1 (15571-1-AP, Proteintech); BCL2 (12789-1-AP, Proteintech); CYCS (A-8, Santa Cruz); GOLGA2 (D6B1, Cell Signaling Technology); PDCD6IP (3A9, Santa Cruz); ITGA3 (21992-1-AP, Proteintech); ANXA1 (AB214486, Abcam); HSP90B1 (9G10, Enzo Life Sciences); CDKN1A (F-5, Santa Cruz); phospho-p53 ser15 (9284, Cell Signaling Technology); GST (rabbit polyclonal, in-house–generated); LRP6 (C47E12, Cell Signaling Technology).

Ab for immunoprecipitation (IP) were directed against: anti-FLAG magnetic Agarose (A36797, Life Technologies); anti-FLAG M2-agarose affinity gel (A2220, Merck Life Science); anti-HA (A2095, Merck Life Science); NUMB (AB21 ^2^); SNX9 (15721-1-AP, Proteintech).

### Engineering of constructs

Expression vectors for p53, MDM2 and NUMB were described in ^1^. The NUMB-1-GFP fusion previously described in ^2^ was recloned into a pLVX-Puro vector (Clontech Laboratories, Inc). To prevent GFP dimerization, the A206K mutation was introduced into the GFP sequence, as described in ^70^. Unless otherwise indicated, when NUMB-FL was transfected into cells, we always refer to NUMB-1.

Mammalian expression vectors for SNX9 were engineered in a pcDNA3 plasmid (Invitrogen, Life Technologies) in frame with a HA or FLAG tag at the C-terminus. GST-SNX9 constructs for bacterial expression were cloned into a pGEX6p-2rbs vector ^71^. The PIP2-YFP(ST) sensor was a gift from Kees Jalink (Addgene plasmid # 170354; http://n2t.net/addgene:170354; RRID:Addgene_170354) ^72^. WT 6XHis-SNX9 pET15b was a gift from Sandra Schmid (Addgene plasmid # 34690; http://n2t.net/addgene:34690; RRID:Addgene_34690) ^73^. The retroviral plasmid H2B-PamCherry was described in ^74^.

To produce the NUMB-GFP11 and SNX9-GFP10 fusion proteins, we first inserted either a synthetic fragment encoding a flexible 25-mer linker-GFP11 or a 30-mer linker-GFP10 into the pcDNA3 vector (Invitrogen, Life Technologies), according to ^43^. The open reading frames (ORFs) of the NUMB-1 and SNX9 genes were subsequently cloned into these vectors at the N-terminus. pCMV GFP1-9 was a gift from Stéphanie Cabantous (Addgene plasmid # 182244; http://n2t.net/addgene:182244; RRID:Addgene_182244) ^43^.

GFP-p53 cDNA was subcloned by PCR from the NUMB-1-GFP plasmid and the p53 plasmid described in ^2^ into the pLVX-Puro vector (Clontech Laboratories, Inc) via NEBuilder HiFi DNA Assembly kit (New England Biolabs) following the manufacturer’s instructions. GFP was placed at the N-terminal position relative to p53.

Site-directed mutagenesis of various constructs was performed using the QuikChange system (Agilent Technologies), following the manufacturer’s protocol. All modifications and inserts were verified by DNA sequencing and/or restriction enzyme digestion.

Throughout the paper, we use HUGO nomenclature for all genes and gene products, except for p53 for which the official symbol, TP53, is much less used in the literature.

### Silencing and KO experiments

Specific siRNAs for total Numb and Numb isoforms and the corresponding control were described previously ^1^. For SNX9 silencing, the siRNA oligonucleotide used was: 5’- UAAGCACUUUGACUGGUUA-3’. Cells were transfected using Lipofectamine RNAiMAX (Invitrogen, Life Technologies) for 72 h (final siRNA concentration, 10 nM). For SNX9 silencing, two consecutive transfection cycles were performed, each lasting 48 h. To create a SNX9 siRNA-resistant construct, silent mutations were introduced into the SNX9 cDNA corresponding to the region recognized by the siRNA oligo, which was therefore mutated from TAAGCACTTTGACTGGTTA to TAAACATTTCGATTGGTTG.

NUMB, SNX9, and p53 KO cell lines were generated via CRISPR/Cas9 genome editing, following the protocol described by Sakuma et al. ^75^. The plasmids pX330A-1x2 (Addgene #58766; RRID:Addgene_58766) and pX330S-2 (Addgene #58778; RRID:Addgene_58778), originally developed by Takashi Yamamoto, were employed for single-guide RNA (sgRNA) expression. The pX330A-1x2 plasmid was modified to include a puromycin resistance cassette downstream of the Cas9 ORF, generating the selection-compatible vector pX330A-1x2 Puro (gift from Diego Pasini).

sgRNAs were designed using the Benchling CRISPR design tool (https://www.benchling.com) and cloned into the aforementioned CRISPR vectors. The selected target sequences were:

5’-GTGACAGATCCAGCGACGAG -3’ (NUMB exon 5),

5’-ATCCTCATGCCATCCCACGC-3’ (NUMB exon 7),

5’-ACCAGCAGCTCCTACACCGG-3’ (p53 exon 4),

5’-TGGGAGAGACCGGCGCACAG-3’ (p53 exon 8),

5’-GAAACATCAAAGGAGAACGA-3’ (SNX9 exon 3),

5’- GGCTGTGGGGAAGTTCACCA-3’ (SNX9 exon 12).

Exon numbering is based on Ensembl transcript reference sequences (p53: ENST00000269305.9; NUMB: ENST00000555238.6; SNX9: ENST00000392185.8). For NUMB, exon numbers exclude the first three untranslated exons. For each gene, dual-sgRNA guides were assembled into the pX330A-1x2 Puro vector via Golden Gate cloning as described by Sakuma, T. et al. ^75^, enabling the simultaneous targeting of two distinct exons. The double NUMB/SNX9-KO in HEK-293 cells was achieved by combining sgRNAs targeting NUMB exon 5 and SNX9 exon 3.

HEK-293 and MCF10A cells were transfected with either the targeting constructs or an empty pX330A-1x2 Puro vector (Ctrl-KO) using jetOPTIMUS (Polyplus) according to the manufacturer’s protocol. At 24 h post-transfection, cells were selected with puromycin (1 μg/mL) for five days and subsequently maintained as a bulk population. For p53-KO, individual clones were isolated by limiting dilution. KO was verified by IF and WB analysis.

NUMB-KO was conducted exclusively in HEK-293 cells to investigate the impact of NUMB depletion on p53 levels in EVs. MCF10A cells were excluded from this analysis because NUMB loss in these cells is known to result in intracellular p53 downregulation, owing to the lack of NUMB-mediated inhibition of MDM2 activity ^1, 2^. This reduction in p53 complicates the interpretation of EV cargo profiles, as it becomes unclear whether altered p53 levels in EVs reflect changes in cargo sorting or merely mirror intracellular depletion. HEK-293 cells were selected to circumvent this confounding effect, as NUMB silencing in these cells does not reduce p53 protein levels, as demonstrated in the present study. This is **likely attributable** to the presence of stably integrated adenoviral E1A and E1B gene products, which modulate p53 dynamics. E1A promotes p53 expression, while E1B inhibits p53-mediated apoptosis and cell cycle arrest, collectively leading to stabilization of p53 protein levels despite NUMB loss ^76, 77^.

HBL-100 p53 silenced cells (Fig. S8B) were obtained by infecting the cells with the shp53 pLKO.1 plasmid (Addgene plasmid # 19119; http://n2t.net/addgene:19119; RRID:Addgene_19119) ^78^. Cells were then selected with puromycin (1 μg/mL) and subsequently maintained as a bulk population.

### RNA purification and quantitative real-time PCR analysis

Total RNA was extracted from cells using the RNeasy Micro Kit (Qiagen) according to the manufacturer’s instructions. Between 500 and 1000 ng of purified RNA, quantified using a NanoDrop spectrophotometer (Thermo Fisher Scientific), were used for first-strand cDNA synthesis with the SuperScript IV VILO Master Mix (Thermo Fisher Scientific) in a final reaction volume of 20 µl.

Quantitative RT-PCR was performed using TaqMan probes (Thermo Fisher Scientific) and TaqMan Universal PCR Master Mix. Each gene was analyzed in at least technical duplicate. The ΔCt method was used to quantify mRNA levels of each target gene, normalized to at least one of the housekeeping genes (GAPDH, GUSB, 18S) from the list below. The 2^-ΔΔCt^ method was used to compare the mRNA levels of each target gene, normalized to the housekeeping genes, relative to an external standard.

Taqman Gene Expression Assay IDs were (Thermo Fisher Scientific): MDM2 (hs00242813_m1), CDKN1A (hs01121172_m1), GAPDH (Hs03929097_g1), GUSB (Hs99999908_m1), 18S (4319413E).

### Recombinant protein expression and purification

GST-fusion proteins were expressed in BL21 Rosetta *E. coli* cells by inducing expression for 6-8 h at 20°C with 0.3 mM IPTG. Cells were lysed in a buffer containing 100 mM Tris-HCl pH 7.6, 300 mM NaCl, 10% glycerol, 0.5 mM EDTA, 1 mM DTT, 0.2 mg/ml lysozyme, and a protease inhibitor cocktail (Complete, EDTA-free, Roche, used here and in all subsequent experiments). Following lysis, the cells were sonicated, and the lysate was clarified by centrifugation at 20,000 g for 30 min. The resulting supernatant was incubated with Glutathione Sepharose 4 Fast-Flow beads (GE Healthcare) to capture the fusion proteins.

For MDM2^216–302^ fragment, the protein was cleaved with PreScission protease overnight at 4°C to remove the GST tag. The cleaved protein was eluted in a desalting buffer (20 mM Tris-HCl, pH 7.6, 40 mM NaCl, 5% glycerol, 1 mM DTT, 0.5 mM EDTA) and then subjected to ion exchange chromatography on a Resource-Q column (GE Healthcare). The MDM2^216–302^ fragment used was a phosphomimetic mutant, in which the known phosphoserine sites were substituted with aspartic acid residues. This mutant, described in ^1^, binds more efficiently to NUMB, likely mimicking the physiological state of unstressed cells ^79^.

For the expression of His-tagged proteins, BL21 Rosetta pLysS *E. coli* cells were induced for 3 h at 30°C with 1 mM IPTG. Cells were lysed in a buffer composed of 100 mM Tris-HCl pH 7.6, 300 mM NaCl, 10% glycerol, 0.2 mg/ml lysozyme, 2 mM 2-mercaptoethanol, 20 mM imidazole, and protease inhibitor cocktail. After sonication, the lysate was clarified by centrifugation at 20,000 g for 30 min. The supernatant was incubated with Ni-NTA beads (QIAGEN), and the bound proteins were eluted with 50 mM Tris-HCl pH 7.6, 150 mM NaCl, and 200 mM imidazole. The eluted proteins were dialyzed overnight at 4°C in a buffer containing 50 mM Tris-HCl pH 7.6, 150 M mNaCl, 0.5 mM EDTA, and 1 mM DTT, with PreScission protease added to remove the His tag, when necessary. The cleaved proteins were subsequently purified using a Superdex-200 column (GE Healthcare) equilibrated with the same buffer. Peak fractions were pooled, concentrated to approximately 10 mg/ml using centrifugal filter concentrators (Amicon), flash frozen in liquid nitrogen, and stored at -80°C.

To purify p53, Expi293 cells were transfected with p53-FLAG and then lysed in JS lysis buffer (50 mM HEPES pH 7.5, 150 mM NaCl, 5 mM EGTA, 1.5 mM MgCl_2_, 10% glycerol, 1% Triton X-100) supplemented with a protease inhibitor cocktail. The cleared lysate was incubated with anti-FLAG M2 agarose affinity beads (Sigma Aldrich) and the retained protein was eluted in 20 mM Tris pH 7.6, 150 mM NaCl, 0.5 mM EDTA, 5% glycerol and 100 µg/ml FLAG peptide (Merck Life Science). After recovery, 1 mM DTT was added and the protein was loaded on a Superose-600 column (GE Healthcare) equilibrated with the same buffer. The peak fractions were pooled and concentrated using centrifugal filters concentrators (Amicon), pre-washed with 0.01% BSA and water, flash frozen in liquid nitrogen, and stored at -80°C.

### Protein studies

#### Immunoblot (IB)

Cells were lysed with ice-cold RIPA buffer (50 mM TRIS pH 7.5, 150 mM NaCl, 1 mM EDTA pH 8, 1% NP-40, 0.5% Na-deoxycholate, 0.1% SDS supplemented with protease inhibitor cocktail) for 30 min in ice. For the experiments in Fig. 7A and Fig. S10B, a phosphatase inhibitor cocktail (PhosSTOP, Roche) was also added. Lysates were cleared by centrifugation at 21,000×g for 30 min at 4°C. Protein concentration was determined using the Pierce BCA protein assay kit (Thermo Fisher Scientific). Lysates were supplemented with 4X Laemmli loading buffer (250 mM TRIS pH 6.8, 40% glycerol, 8% SDS, 0.02% bromophenol blue, 200 mM DTT), boiled for 10 min at 95°C, and loaded onto pre-cast gels (Biorad or Thermo Fisher Scientific) before being transferred to nitrocellulose membranes using a Trans-Blot system (Biorad), according to the manufacturer’s guidelines. The membranes were then blocked with 5% milk in TBS containing 0.1% Tween20 (TBS-T), followed by incubation with the primary antibody according to the manufacturer’s instructions. After three washes with TBS-T, the membranes were incubated with the appropriate horseradish peroxidase-conjugated secondary antibody. Following three additional washes, the signal was visualized by developing films (Amersham) or using the Invitrogen iBright Imaging System with ECL detection reagents from Amersham or Biorad.

In several IB and co-IP experiments, samples were run on separate gels to detect proteins with similar molecular weights. Gels were then sectioned in the regions of interest to enable probing with different antibodies. In some instances, membranes were also stripped and reprobed. All immunoblots were independently repeated at least twice.

#### Co-Immunoprecipitation (o-IP)

Cells were lysed in ice-cold JS buffer for co-IP experiments (50 mM HEPES pH 7.5, 150 mM NaCl, 5 mM EGTA, 1.5 mM MgCl_2_, 10% glycerol, 1% Triton X-100 supplemented with a protease inhibitor cocktail) for 30 min in ice. Lysates were cleared by centrifugation at 21,000 ×g for 30 min at 4°C. Protein concentration was determined using the Bradford assay (Biorad). An aliquot of cellular lysate was directly supplemented with 4X Laemmli loading buffer for loading as total lysate. At least 1 mg of cellular lysate was incubated with FLAG beads (anti-Flag M2-agarose affinity gel, Sigma-Aldrich, or anti-FLAG magnetic Agarose, Life Technologies) or HA beads (anti-HA Agarose, Merck Life Science), using approximately 6-8 μL of beads slurry per 1 mg of lysate.

For the experiment in Fig. 6B, lysates of EVs, purified by ultracentrifugation from 6 x 10^7^ SNX9-FLAG expressing Expi293 cells, were incubated with 10 μl anti-FLAG magnetic agarose beads.

For immunoprecipitation of endogenous NUMB or SNX9, the corresponding antibody (∼ 2 μg Ab/1 mg lysate) was pre-conjugated to Protein A/G magnetic beads (Pierce) before being added to at least 3 mg of lysate, previously pre-cleared with empty beads.

In control samples, a corresponding peptide was added to the lysate at a final concentration of 100 µg/mL. The beads were incubated for 2 h at 4°C with agitation, then washed three times in JS buffer. Finally, the protein complexes were eluted in 2X Laemmli buffer.

In all corresponding IB, the lysate lane is ∼1/50^th^ of the IP, with the exception of Fig. 1C, 1E, 1G, where the lysate lane is ∼1/150^th^ of the IP.

#### In vitro binding assays with purified proteins (Figs. 2C, 4D, S2B, S4A-B, S5D, S6D)

GST-tagged proteins (1 µM) were immobilized onto GSH beads (GE Healthcare) and incubated for 2 h at 4°C with the indicated proteins, at the concentrations specified in the text, in 10 mM Hepes pH 7.5, 150 mM NaCl, 5% glycerol, and 0.1% Tween-20 (final volume 200 µl), with the exception of Fig. 4D in which the proteins concentrations used are specified in the figure legend and Fig. S6D where the GST-tagged proteins were at a concentration of 0.5 µM and SNX9 proteins at a concentration of 1.5 µM. After washing, the protein complexes were eluted in 2X Laemmli buffer and analyzed by SDS-PAGE. Detection was with Coomassie. In these gels, half of each sample was loaded (with the exception of Fig. 4D where 1/20th was loaded); the input lanes contained 500 ng of the corresponding protein used for incubation with the GSH beads, with the exception of Fig. S6D where the input lane contained 1000 ng.

#### Pull-down assays with cellular lysates (Fig. S4C-E)

GST-tagged proteins (0.1 µM) were immobilized onto GSH beads and incubated for 2 h at 4°C with 1 mg of HEK-293 cellular lysate prepared in JS buffer. After washing, the proteins retained on the beads were eluted in 2X Laemmli buffer, separated by SDS-PAGE and detected by IB. In these experiments, the “lysate” lanes were 1/30^th^ of the loaded pull-down fraction.

#### Analytical size exclusion chromatography (SEC)

For the SEC analyses presented in Figs. S3A,C, S6C, and S7E, all proteins were combined to a final concentration of 100 µM (in the case of SNX9 FL, the concentration was calculated considering a dimeric molecule), except for MDM2, which was used at 400 µM. The samples were loaded onto a Superdex 200 Increase 5/150 GL (Cytiva) column equilibrated in 20 mM Tris pH 7.6, 200 mM NaCl, 0.5 mM EDTA pH 8, and 1 mM DTT. Eluted species were monitored by absorbance at 280 nm and subsequently analyzed by SDS-PAGE followed by Coomassie staining, loading equal quantities of the fractions.

### Nanoparticle Tracking Analysis and Tunable Resistive Pulse Sensing (TRPS)

For the experiment shown in Fig. S9D, the mean EV concentration and diameter were analyzed by TRPS using an Exoid instrument (iZON Science Ltd., Christchurch, New Zealand). TRPS measurements were performed with a nanopore NP200 (iZON Science Ltd.). The nanopore stretch was 43.5-45 nm, the applied current ranged from 120-140 nA, and the injection pressure ranged from 800-2400 Pa. The stability of the baseline current (i.e., background noise) was always below 10 pA, in accordance with the manufacturer’s instructions for optimal measurements. Samples were diluted in PBS to achieve a blockade rate between 100–1500 particles. A total of 500 particles were recorded per pressure; when the particle rate was below 100, each pressure was recorded for 2 min.

For the experiment shown in Fig. S9E, nanoparticle tracking analysis was conducted using a NanoSight NS300 system (NanoSight NTA 2.3, Salisbury, UK) in accordance with the manufacturer’s guidelines. Each sample was prepared diluted in PBS, and analyzed under consistent post-acquisition settings optimized prior to measurement. For each sample, three 60-second videos were captured and processed to determine particle concentration.

### Transmission Electron Microscopy

For negative stain grids preparation, liposomes were diluted to 10 µM in 50 mM Hepes pH 7.5, 1 mM DTT, 1 mM EDTA pH 8.0, 180 mM NaCl and 90 mM saccharose, while purified EVs derived from HEK-293 cells were fixed in PBS with 2% PFA for 5 min at RT. Four µl of sample were applied to freshly glow-discharged carbon film only on Cu 400 mesh grids (Ted Pella) and incubated for 1 min. Excess of sample was blotted away with filter paper and the grids were stained for 1 min with 10 µl of 2% (w/v) uranyl acetate. Grids were then blotted, dried and visualized using a 120 kV JEOL JEM 1200-EX II equipped with TEM CCD camera Mega View III (Olympus). Images were recorded at a nominal magnification of 50,000-100,000X.

### Mass spectrometry

Entire SDS-PAGE gel lanes were excised and finely chopped into small fragments. Disulfide bonds within the proteins were reduced using 10 mM dithiothreitol (DTT) in 50 mM ammonium bicarbonate (NH4HCO3) at 56°C for 55 minutes, followed by alkylation with 55 mM iodoacetamide in the same buffer for 20 minutes at room temperature. Protein digestion was carried out using trypsin (0.1 μg/μL in 50 mM NH4HCO3) with incubation at 37°C overnight. Peptides were subsequently extracted from the gel pieces using a solution of 95% acetonitrile and 5% formic acid. The eluates were dried and desalted using the StageTip method^80^. Peptide samples were analyzed using a quadrupole Orbitrap Q Exactive HF mass spectrometer (Thermo Scientific). Separation of peptides was performed on a UHPLC Easy-nLC 1000 system (Thermo Scientific) with a linear gradient from 95% solvent A (2% acetonitrile, 0.1% formic acid) to 12% solvent B (80% acetonitrile, 0.1% formic acid) in 3 minutes, then to 50% solvent B over 33 minutes, followed by an increase to 100% solvent B within 2 minutes, maintained for further 3 minutes. The flow rate was maintained at 0.25 μL/min. The LC system was interfaced with a 23-cm fused silica emitter (75 μm internal diameter; New Objective, Woburn, MA, USA), which was packed in-house with ReproSil-Pur C18-AQ 1.9 μm particles (Dr. Maisch GmbH, Ammerbuch, Germany) using a high-pressure packing device (Proxeon, Odense, Denmark). The mass spectrometer was operated in positive data-dependent acquisition (DDA) mode ^81^. Dynamic exclusion was active with a 15-second exclusion window. Full MS1 spectra were acquired at a resolution of 60,000 with an automatic gain control (AGC) target of 3 × 106 and a maximum injection time of 20 ms. MS2 scans were recorded at a resolution of 15,000, using an AGC target of 1 × 10^5^, with the injection time of 80 ms and a normalized collision energy (NCE) of 28. Each acquisition cycle consisted of one full MS1 scan (range: 300–1,650 m/z) followed by 15 MS2 scans using a 1.2 m/z isolation window. All raw data were processed using MaxQuant V1.5.2.8 PMID: 19029910, searched against the Uniprot-Complete Proteome_HomoSapiens_2017_database. Carbamidomethylation of cysteine residues was defined as a fixed modification, while oxidation of methionine and acetylation of protein N-termini were treated as variable modifications.

The mass spectrometry proteomics data have been deposited to the ProteomeXchange Consortium via the PRIDE partner repository PMID: 39494541 with the dataset identifier PXD064868.

### Mass photometry analysis

Purified recombinant full-length SNX9 was analyzed by mass photometry with a Two^MP^ instrument (Refeyn). The protein sample was diluted to 10 nM in 20 mM Tris pH 7.6, 200 mM NaCl, 0.5 mM EDTA and 1 mM DTT. The instrument was calibrated using a solution containing a mix of bovine thyroglobulin (Sigma-Aldrich), equine apoferritin (Sigma-Aldrich) and bovine serum albumin (G-Biosciences).

### Intracellular ATP levels detection

Intracellular ATP levels were measured as a surrogate marker to indirectly assess cell proliferation, using the CellTiter-Glo assay (Promega) according to the manufacturer’s instructions. SAOS2 cells were plated in white-walled 96-well plates and treated with EVs. Following treatment, cells were incubated for 10 min at RT with gentle agitation in CellTiter-Glo Reagent, which contains a luciferase substrate in a stabilized buffer. Luminescence, proportional to ATP concentration, was then recorded using a GloMax luminometer. The mean ± SD of ten wells per condition from two independent experiments is shown.

### Imaging studies

#### Immunofluorescence (IF)

Cells were seeded on coverslips pre-treated with fibronectin (Roche) at a final concentration of 5 µg/cm^2^, when required, or directly in tissue culture-treated chambered coverslips (Ibidi). For the experiments shown in Fig. 7C and Figs. S10C and S10E, cells were treated with an acid wash (100 mM Glycine-HCl, pH 2.2) prior to fixation, to remove potential EVs attached to the PM and not internalized.

Cells were washed with PBS and fixed with 4% PFA for 12 min at RT, then washed twice with PBS and permeabilized with 0.1% Triton X-100 in PBS for 10 min, or with 0.005% saponin, where indicated, to reduce nuclear staining (in this case, the same concentration of saponin was maintained in all subsequent steps). Blocking was performed with 10% normal donkey serum (Jackson ImmunoResearch) in PBS for 30 min. Cells were washed twice with 1% BSA in PBS and incubated with primary antibodies diluted in 1% BSA in PBS for 1 h at RT in a humid chamber. After washing cells were incubated with appropriate secondary antibodies diluted in 1% BSA in PBS for 1 h in a dark, humid chamber. After washing, nuclei were stained with DAPI and cells were washed twice with PBS, briefly rinsed in water, and mounted on glass slides using Mowiol mounting medium (Sigma-Aldrich), then dried at RT in the dark. Cells plated into the chambered coverslips were conserved in PBS at 4°C after DAPI staining.

Images were obtained using a Leica TCS SP8 confocal microscope equipped with a 63X oil objective unless otherwise indicated and processed using tools available via ImageJ/Fiji software (National Institutes of Health) and Photoshop (Adobe). To improve visual clarity for publication, where required, brightness and contrast were adjusted equally across all images, ensuring that the adjustments did not affect the scientific interpretation and no data were altered or obscured.

#### Time-lapse experiments

For the time-lapse experiments shown in Fig. S7C, MCF10A cells were plated in glass-bottom 35 mm dishes and transfected as indicated. After 18 h, the PM was stained using CellMask Plasma Membrane Stain Orange (Thermo Fisher Scientific), according to the manufacturer’s instructions, and the cells were then imaged by time-lapse video microscopy using Leica Confocal SP8 equipped with a Leica DMi8 inverted microscope with PMTs and hybrid detectors, a water immersion objective 20X PL APO (NA 0.75) and an incubation chamber (Evotec/Okolab) to maintain the cells at 37°C and 5% of CO_2_. Images were acquired every ∼30 sec until the end of the treatment.

For the quantification of the green signal at the PM, analysis was performed using ImageJ software. For each cell, five random regions of approximately 2 μm at the PM level were selected at both the initial and final time points. The ratio of the mean green intensity in these regions between the initial and final time points was then calculated for each cell.

#### dSTORM microscopy

Single-molecule imaging of epithelial HBL-100 cells was performed with a commercial inverted Nikon Eclipse Ti2 microscope (Nikon instruments, Tokyo, Japan), equipped with a super-resolution Nikon N-STORM module. The system was configured for either total internal reflection fluorescence (TIRF) or oblique incidence excitation, employing a continuous activation mode (both the activator and imaging lasers are continuously on). The fully motorized automated microscope was controlled by the NIS Elements software (version 5.42.07). Alexa Fluor 647 and Cy3 dyes were excited by using 561 nm (240 mW) and 647 nm (360 mW) laser wavelengths respectively (L4Cc combiner, Oxxius S.A., Lannion, France), while a 405-nm solid-state laser (216 mW) was used for activation of dyes. A multi-band dichroic mirror (C-NSTORM QUAD 405/488/561/647 FILTER SET; Chroma), combined with 561 nm- and 647 nm-filter cubes (IDEX Health & Science, Semrock Brightline®, Rochester, NY, USA), was used to filter the fluorescence excitation. Emission light was filtered by the Optospin Filter Wheel (Cairn Research Ltd, Faversham, Kent UK). The fluorescence emission of all channels was collected through a Nikon CFI SR Apochromat TIRF 100× oil objective (1.49 NA) and finally detected by the digital CMOS camera (Dual ORCA Flash 4.0 Digital CMOS camera C13440, Hamamatsu, Tokyo, Japan). The number of frames and exposure time per channel depends by the density pattern of the immunostaining and the dye blinking state (15,000 frames in continuous mode at 30 ms of exposure time per channel). Z drift was minimized thanks to a hardware autofocusing system (Perfect Focus System (PFS) (Nikon instruments, Tokyo, Japan). Under constant illumination and buffer conditions, the dyes typically started in the fluorescent state, switched to a dark state and spontaneously recovered to a fluorescent state several times before photobleaching. The power of both activation and excitation lasers is dependent on the blinking efficiency of the single fluorophores. The modulation of lasers was adjusted with neutral density filters to reduce the photobleaching effect during the preliminary observation for the optimization of the acquisition parameters.

#### dSTORM imaging buffer

Single-molecule dSTORM imaging for each dye was performed in an imaging buffer containing 10 mM Tris (pH 8.0), 50 mM NaCl, 10% Glucose, an oxygen-scavenging system “GLOX” (5.6 mg/mL Glucose Oxidase, 0.34 mg/mL Catalase) and 100 mM mercaptoethylamine. The solution was kept at -20°C and used within 2-3 weeks from preparation.

#### TIRF microscopy

Images were acquired by a Nikon CFI SR Apochromat TIRF 100× oil objective (1.49 NA), configured for total internal reflection imaging, following the light-path explained above (“dSTORM microscopy” section). The critical angle for TIRF illumination was set with the H-TIRF module, integrated in NIS Elements software. The multi-parameters acquisition was performed using a LU-NV laser unit (Nikon instruments, Tokyo, Japan) equipped with 5 laser lines: 405 nm (23.1 mW), 440 nm (25.5 mW) 488 nm (79.1 mW), 561 nm (79 mW), 647 nm (137 mW). Image format was set to 1024×1024 pixels, pixel size of ∼0.06 μm, to optimize homogeneity of illumination in total internal reflection conditions.

#### dSTORM image reconstruction and pixel-based colocalization analysis

Single molecule localization fitting was performed with Offline N-STORM Analysis module (NIS Elements software, version 5.42.07) considering drift and chromatic aberrations. Before exporting STORM images, the Gaussian size of the single localizations and the format of the reconstructed images were set. In the super-resolution image reconstruction, each molecule is represented by a Gaussian spot localized by the centroid position, with the localization precision obtained from the single-molecule fit, and by an amplitude value related to the number of emitted photons. We finally generated a dual-color STORM image of 41 × 41 μm (10 nm/px) with a Gaussian size of 10 and 50 nm. A 50 nm-width PSF can efficiently represent the molecular tree formed by the primary and secondary antibody complex on the target protein avoiding the degeneration introduced by multiple emissions.

Colocalization analysis was performed according to an Image Cross-Correlation Spectroscopy pipeline ^82^. Briefly, images cross-correlation and auto-correlation functions were obtained by averaging over the pixels contained in a selected region of interest (e.g., PM). A mask of the PM was created from the spatial distribution of target proteins. The mask covered an area of ∼2 μm from the cell periphery. The amplitude parameters from the resulting curves were employed to calculate two coefficients of localizations whose arithmetic mean provides the colocalizing fraction *f_ICCS_*. The measured widths are instead related to the broadening of the cross-correlation function that is a parameter sensitive only to the distance *d* between correlated particles ^83^.

#### Object-based colocalization analysis of TIRF data

TIRF images were analyzed by A.M.I.CO. (Automated Microscopy for Image CytOmetry) analysis package developed within the open-source ImageJ platform using its macro programming language. The package was described in detail in the previous works ^84^. For every parameter, the quantification of colocalization rate was performed by the distance analysis module integrated in A.M.I.CO. Briefly, the three channels were segmented to identify the object of interest (fluorescent spots) and calculate the geometrical centroid of the object. The degree of co-localization was calculated depending on TIRF optical resolution and the size of objects investigated. In this approach, a spherical region with a radius of 300 nm (5 px) was defined around the center of the object, considered as the cut-off distance for proximity. Colocalization was determined if the center of another object fell within this region.

### Liposome preparation and liposome binding assays

Unilamellar liposomes were prepared by combining 90% phosphatidylcholine (PC), 5% phosphatidylserine (PS), and 5% of either phosphatidylinositol (4,5)-bisphosphate (PI(4,5)P₂) or phosphatidylinositol (4)-phosphate (PI(4)P). The lipid mixture was dried under a stream of nitrogen for 1 h. Dried lipids were then resuspended in buffer containing 50 mM HEPES pH 7.5, 100 mM NaCl, 1 mM EDTA, 1 mM DTT, and 180 mM sucrose to a final lipid concentration of 6.5 mM. Liposomes were subsequently extruded using a Micro Extruder set (Avanti Polar Lipids) equipped with polycarbonate membranes with a pore size of 200 nm. The extruded liposomes were stored at 4°C to preserve stability.

For binding assays, liposomes were incubated with recombinant proteins at the concentrations indicated in the legend to Fig. 4, at a final liposome concentration of 3.25 mM, in a buffer containing (after mixing proteins and liposomes) 50 mM HEPES pH 7.5, 180 mM NaCl, 1 mM EDTA, 1 mM DTT, and 90 mM sucrose. After a 10 min incubation at RT, samples were ultracentrifuged at 213,000 × g for 1 h at 4°C. Pellets were resuspended in 2× Laemmli buffer, and equal amounts of total, supernatant, and pellet fractions were subsequently analyzed by SDS-PAGE, followed by Coomassie staining, except for Fig. 4C, where the pellet fraction was loaded in a 3-fold excess relative to the total fraction.

### Isolation of EVs and uptake experiments

Cell lines were grown for 24 h in their respective media supplemented with 2% EV-depleted serum, except for Expi293 cells, which do not require serum. EV-depleted serum was prepared by ultracentrifugation at 150,000 × g for 18 h at 4 °C, followed by filtration through a 0.22 μm filter. The conditioned medium was collected into 50 ml tubes and processed through a series of centrifugation steps at 4 °C: 1,200 rpm for 5 min, 3,000 rpm for 10 min, and 4,000 rpm for 30 min. The supernatant was then filtered through a 0.45 μm membrane and concentrated ∼20-fold using 100 kDa molecular weight cutoff centrifugal filters (Amicon). The concentrate was further centrifuged at 10,000 × g for 30 min at 4 °C and the resulting supernatant was transferred into pre-chilled ultracentrifuge tubes (previously rinsed with 70% ethanol) and subjected to ultracentrifugation at 110,000 × g for 2 h at 4 °C. EV pellets were washed in PBS and ultracentrifuged again at 110,000 × g for 1.5 h. Final EV pellets were lysed in RIPA buffer supplemented with protease inhibitors.

For SDS-PAGE analysis, 1 μg of total cell lysate and 2 μg of purified EVs were loaded per lane, except in Figs. 6C and 6F, where 1 μg of total cell lysate and 1/5^th^ of the EV samples were loaded for each condition.

In Figs. 7C,D,G and S10C,E, EVs were isolated via ultrafiltration, omitting the ultracentrifugation steps to preserve EV integrity and enhance uptake, as described ^85^. To prevent exosome aggregation, 25 mM trehalose was added to the medium before ultrafiltration, following published protocols ^86^. Purified EVs were applied for 8 h to recipient cells, in medium containing EVs-depleted serum, at an approximate donor-to-recipient cell ratio of 100:1, consistent with previous reports indicating the low efficiency of EV uptake ^87^. In Fig. 7G, EVs were added every 24 h to SAOS2 recipient cells and medium and EVs were replenished daily. In Fig. S10E, EVs were added to SAOS2 cells for 15 h before IF analysis. In Fig. S9E, MCF10A cells were cultured in serum-free medium for 20 min to minimize the uptake of newly released EVs. EVs were purified via ultrafiltration to maximize their recovery, which were otherwise partially lost during ultracentrifugation.

In Fig. 7F, MCF10A donor cells (Ctrl KO, SNX9-KO, and p53-KO) were seeded in 15 cm dishes. In parallel, MCF10A p53 KO H2B-Cherry recipient cells were plated on coverslips in separate dishes. Seeding densities were adjusted to reach a donor-to-recipient cell ratio of ∼50:1. The next day, medium was replaced with complete medium containing 4% EV-depleted serum, using the minimum volume necessary. Coverslips were transferred into dishes containing donor cells, allowing them to float in the medium. After 48 h, where indicated, cells were treated with 50 μM etoposide for 8 h. Coverslips were then retrieved, washed extensively with PBS, and processed for RNA extraction.

In Fig. S9B, EV pellets purified by ultracentrifugation were resuspended in PBS and divided into three aliquots. Where indicated, samples were treated with 5 μg/ml proteinase K and/or 1% Triton X-100 and incubated at 37 °C for 30 min. Reactions were stopped by adding a protease inhibitor cocktail and 1 mM PMSF, followed by a 10 min incubation on ice. Samples were then lysed by adding Laemmli buffer.

### Statistical analysis

All statistical analyses were performed using GraphPad Prism. For the analysis in Fig. 3D, a Fisher’s exact test was used to compare the distribution of positive cells between the two groups. One-way ANOVA was performed for Figs. 7G and S9D. Data from Figs. 5A, 6E, 7D, 7F, S5C, and S7C were analyzed using an unpaired parametric t-test. For Figs. 7B and S10B, a paired parametric t-test was used.

## Supporting information

merged file containing all supplementary figures

## ACKNOWLEDGMENTS

We thank Massimo Boiocchi of Centro Grandi Strumenti Electron Microscopy TEM Facility at University of Pavia for his support and assistance in this work. We thank the IEO imaging and Cell Culture Units.

## Funding

Associazione Italiana per la Ricerca sul Cancro (AIRC): IG #22811 to LL, IG #24415 to SS, IG #27014 to GP, IG #31110 to SP, IG #23060 to PPDF.

European Research Council: ERC-CoG2020 #101002280 to SS.

The Italian Ministry of University and Scientific Research: PRIN 2022 Prot. 2022W93FTW to SS, PRIN 2020 Prot. 2020R2BP2E to PPDF, Next Generation EU-CN00000041-National Center for Gene Therapy and Drugs based on RNA Technology to PPDF

The Italian Ministry of Health: Ricerca Corrente 2023-2024 and 5 per 1000 funds, RF-2021-12373957 to PPDF, PNRR-MCNT2-2023-12378490 to PPDF

Roberta Cacciatore is supported by an AIRC fellowship for Italy.

## Author contributions

Conceptualization: PPDF, INC

Methodology: LS, AC, AB, SS, LL

Data Analysis: SF, LL, INC, PPDF

Investigation: RC, AB, IS-L, GC, CRZ, SR, SPi, EZ, CZ, VM

Funding acquisition: PPDF, LL, SS, SPe, GP

Supervision: MF, GP, AB, SPe, SS, INC, PPDF

Writing – original draft: PPDF, INC, RHG

Writing – review & editing: All authors

## REFERENCES

1. Colaluca, I.N., Basile, A., Freiburger, L., D’Uva, V., Disalvatore, D., Vecchi, M., Confalonieri, S., Tosoni, D., Cecatiello, V., Malabarba, M.G., et al. (2018). A Numb-Mdm2 fuzzy complex reveals an isoform-specific involvement of Numb in breast cancer. J Cell Biol 217, 745–762. 10.1083/jcb.201709092

2. Colaluca, I.N., Tosoni, D., Nuciforo, P., Senic-Matuglia, F., Galimberti, V., Viale, G., Pece, S., and Di Fiore, P.P. (2008). NUMB controls p53 tumour suppressor activity. Nature 451, 76–80. 10.1038/nature06412

3. Guo, Y., Zhang, K., Cheng, C., Ji, Z., Wang, X., Wang, M., Chu, M., Tang, D.G., Zhu, H.H., and Gao, W.Q. (2017). Numb(-/low) Enriches a Castration-Resistant Prostate Cancer Cell Subpopulation Associated with Enhanced Notch and Hedgehog Signaling. Clin Cancer Res 23, 6744–6756. 10.1158/1078-0432.CCR-17-0913

4. Xu, T., Verhagen, M., Joosten, R., Sun, W., Sacchetti, A., Munoz Sagredo, L., Orian-Rousseau, V., and Fodde, R. (2022). Alternative splicing downstream of EMT enhances phenotypic plasticity and malignant behavior in colon cancer. Elife 11. 10.7554/eLife.82006

5. Filippone, M.G., Freddi, S., Zecchini, S., Restelli, S., Colaluca, I.N., Bertalot, G., Pece, S., Tosoni, D., and Di Fiore, P.P. (2022). Aberrant phosphorylation inactivates Numb in breast cancer causing expansion of the stem cell pool. J Cell Biol 221. 10.1083/jcb.202112001

6. Shu, Y., Xu, Q., Xu, Y., Tao, Q., Shao, M., Cao, X., Chen, Y., Wu, Z., Chen, M., Zhou, Y., et al. (2021). Loss of Numb promotes hepatic progenitor expansion and intrahepatic cholangiocarcinoma by enhancing Notch signaling. Cell Death Dis 12, 966. 10.1038/s41419-021-04263-w

7. Wang, X., Cai, J., Zhao, L., Zhang, D., Xu, G., Hu, J., Zhang, T., and Jin, M. (2021). NUMB suppression by miR-9-5P enhances CD44(+) prostate cancer stem cell growth and metastasis. Sci Rep 11, 11210. 10.1038/s41598-021-90700-x

8. Tosoni, D., Pambianco, S., Ekalle Soppo, B., Zecchini, S., Bertalot, G., Pruneri, G., Viale, G., Di Fiore, P.P., and Pece, S. (2017). Pre-clinical validation of a selective anti-cancer stem cell therapy for Numb-deficient human breast cancers. EMBO Mol Med 9, 655–671. 10.15252/emmm.201606940

9. Sheng, W., Dong, M., Chen, C., Wang, Z., Li, Y., Wang, K., Li, Y., and Zhou, J. (2017). Cooperation of Musashi-2, Numb, MDM2, and P53 in drug resistance and malignant biology of pancreatic cancer. FASEB J 31, 2429–2438. 10.1096/fj.201601240R

10. Liu, D., Cui, L., Wang, Y., Yang, G., He, J., Hao, R., Fan, C., Qu, M., Liu, Z., Wang, M., et al. (2016). Hepatitis B e antigen and its precursors promote the progress of hepatocellular carcinoma by interacting with NUMB and decreasing p53 activity. Hepatology 64, 390–404. 10.1002/hep.28594

11. Liu, C., Liu, L., Chen, X., Cheng, J., Zhang, H., Shen, J., Shan, J., Xu, Y., Yang, Z., Lai, M., and Qian, C. (2016). Sox9 regulates self-renewal and tumorigenicity by promoting symmetrical cell division of cancer stem cells in hepatocellular carcinoma. Hepatology 64, 117–129. 10.1002/hep.28509

12. Bu, P., Wang, L., Chen, K.Y., Srinivasan, T., Murthy, P.K., Tung, K.L., Varanko, A.K., Chen, H.J., Ai, Y., King, S., et al. (2016). A miR-34a-Numb Feedforward Loop Triggered by Inflammation Regulates Asymmetric Stem Cell Division in Intestine and Colon Cancer. Cell Stem Cell 18, 189–202. 10.1016/j.stem.2016.01.006

13. Tosoni, D., Zecchini, S., Coazzoli, M., Colaluca, I., Mazzarol, G., Rubio, A., Caccia, M., Villa, E., Zilian, O., Di Fiore, P.P., and Pece, S. (2015). The Numb/p53 circuitry couples replicative self-renewal and tumor suppression in mammary epithelial cells. Journal of Cell Biology 211, 845–862. 10.1083/jcb.201505037

14. Siddique, H.R., Feldman, D.E., Chen, C.L., Punj, V., Tokumitsu, H., and Machida, K. (2015). NUMB phosphorylation destabilizes p53 and promotes self-renewal of tumor-initiating cells by a NANOG-dependent mechanism in liver cancer. Hepatology 62, 1466–1479. 10.1002/hep.27987

15. Forloni, M., Dogra, S.K., Dong, Y., Conte, D., Jr., Ou, J., Zhu, L.J., Deng, A., Mahalingam, M., Green, M.R., and Wajapeyee, N. (2014). miR-146a promotes the initiation and progression of melanoma by activating Notch signaling. Elife 3, e01460. 10.7554/eLife.01460

16. Ito, T., Kwon, H.Y., Zimdahl, B., Congdon, K.L., Blum, J., Lento, W.E., Zhao, C., Lagoo, A., Gerrard, G., Foroni, L., et al. (2010). Regulation of myeloid leukaemia by the cell-fate determinant Musashi. Nature 466, 765–768. 10.1038/nature09171

17. Westhoff, B., Colaluca, I.N., D’Ario, G., Donzelli, M., Tosoni, D., Volorio, S., Pelosi, G., Spaggiari, L., Mazzarol, G., Viale, G., et al. (2009). Alterations of the Notch pathway in lung cancer. P Natl Acad Sci USA 106, 22293–22298. 10.1073/pnas.0907781106

18. Pece, S., Serresi, M., Santolini, E., Capra, M., Hulleman, E., Galimberti, V., Zurrida, S., Maisonneuve, P., Viale, G., and Di Fiore, P.P. (2004). Loss of negative regulation by Numb over Notch is relevant to human breast carcinogenesis. J Cell Biol 167, 215–221. 10.1083/jcb.200406140

19. Rhyu, M.S., Jan, L.Y., and Jan, Y.N. (1994). Asymmetric distribution of numb protein during division of the sensory organ precursor cell confers distinct fates to daughter cells. Cell 76, 477–491. 10.1016/0092-8674(94)90112-0

20. Spana, E.P., Kopczynski, C., Goodman, C.S., and Doe, C.Q. (1995). Asymmetric Localization of Numb Autonomously Determines Sibling Neuron Identity in the Drosophila Cns. Development 121, 3489–3494. 10.1242/dev.121.11.3489

21. Uemura, T., Shepherd, S., Ackerman, L., Jan, L.Y., and Jan, Y.N. (1989). numb, a gene required in determination of cell fate during sensory organ formation in Drosophila embryos. Cell 58, 349–360. 10.1016/0092-8674(89)90849-0

22. Caussinus, E., and Gonzalez, C. (2005). Induction of tumor growth by altered stem-cell asymmetric division in. Nature Genetics 37, 1125–1129. 10.1038/ng1632

23. Babaoglan, A.B., O’Connor-Giles, K.M., Mistry, H., Schickedanz, A., Wilson, B.A., and Skeath, J.B. (2009). Sanpodo: a context-dependent activator and inhibitor of Notch signaling during asymmetric divisions. Development 136, 4089–4098. 10.1242/dev.040386

24. Frise, E., Knoblich, J.A., YoungerShepherd, S., Jan, L.Y., and Jan, Y.N. (1996). The Drosophila Numb protein inhibits signaling of the Notch receptor during cell-cell interaction in sensory organ lineage. P Natl Acad Sci USA 93, 11925–11932. 10.1073/pnas.93.21.11925

25. Guo, M., Jan, L.Y., and Jan, Y.N. (1996). Control of daughter cell fates during asymmetric division: Interaction of numb and notch. Neuron 17, 27–41. 10.1016/S0896-6273(00)80278-0

26. Spana, E.P., and Doe, C.Q. (1996). Numb antagonizes notch signaling to specify sibling neuron cell fates. Neuron 17, 21–26. 10.1016/S0896-6273(00)80277-9

27. Confalonieri, S., Colaluca, I.N., Basile, A., Pece, S., and Di Fiore, P.P. (2019). Exon 3 of the NUMB Gene Emerged in the Chordate Lineage Coopting the NUMB Protein to the Regulation of MDM2. G3 (Bethesda) 9, 3359–3367. 10.1534/g3.119.400494

28. Cotton, M., Benhra, N., and Le Borgne, R. (2013). Numb inhibits the recycling of Sanpodo in Drosophila sensory organ precursor. Curr Biol 23, 581–587. 10.1016/j.cub.2013.02.020

29. Berdnik, D., Török, T., González-Gaitán, M., and Knoblich, J.A. (2002). The endocytic protein α-adaptin is required for numb-mediated asymmetric cell division in. Developmental Cell 3, 221–231. 10.1016/S1534-5807(02)00215-0

30. Langevin, J., Le Borgne, R., Rosenfeld, F., Gho, M., Schweisguth, F., and Bellaiche, Y. (2005). Lethal giant larvae controls the localization of notch-signaling regulators numb, neuralized, and Sanpodo in Drosophila sensory-organ precursor cells. Curr Biol 15, 955–962. 10.1016/j.cub.2005.04.054

31. Hutterer, A., and Knoblich, J.A. (2005). Numb and α-adaptin regulate Sanpodo endocytosis to specify cell fate in external sensory organs. Embo Reports 6, 836–842. 10.1038/sj.embor.7400500

32. McGill, M.A., Dho, S.E., Weinmaster, G., and McGlade, C.J. (2009). Numb regulates post-endocytic trafficking and degradation of Notch1. J Biol Chem 284, 26427–26438. 10.1074/jbc.M109.014845

33. Santolini, E., Puri, C., Salcini, A.E., Gagliani, M.C., Pelicci, P.G., Tacchetti, C., and Di Fiore, P.P. (2000). Numb is an endocytic protein. Journal of Cell Biology 151, 1345–1351. 10.1083/jcb.151.6.1345

34. Nilsson, L., Conradt, B., Ruaud, A.F., Chen, C.C.H., Hatzold, J., Bessereau, J.L., Grant, B.D., and Tuck, S. (2008). negatively regulates endocytic recycling. Genetics 179, 375–387. 10.1534/genetics.108.087247

35. Nilsson, L., Jonsson, E., and Tuck, S. (2011). Caenorhabditis elegans numb inhibits endocytic recycling by binding TAT-1 aminophospholipid translocase. Traffic 12, 1839–1849. 10.1111/j.1600-0854.2011.01271.x

36. Sabbioni, S., Filippone, M.G., Amadori, L., Confalonieri, S., Bonfanti, R., Capoano, S., Colaluca, I.N., Freddi, S., Bertalot, G., Faga, G., et al. (2025). The CRL7(FBXW8) Complex Controls the Mammary Stem Cell Compartment through Regulation of NUMB Levels. Adv Sci (Weinh), e2405812. 10.1002/advs.202405812

37. Dho, S.E., Othman, K., Zhang, Y., and McGlade, C.J. (2025). NUMB alternative splicing and isoform-specific functions in development and disease. J Biol Chem 301, 108215. 10.1016/j.jbc.2025.108215

38. Gazave, E., Lapebie, P., Richards, G.S., Brunet, F., Ereskovsky, A.V., Degnan, B.M., Borchiellini, C., Vervoort, M., and Renard, E. (2009). Origin and evolution of the Notch signalling pathway: an overview from eukaryotic genomes. BMC Evol Biol 9, 249. 10.1186/1471-2148-9-249

39. Bendris, N., and Schmid, S.L. (2017). Endocytosis, Metastasis and Beyond: Multiple Facets of SNX9. Trends Cell Biol 27, 189–200. 10.1016/j.tcb.2016.11.001

40. Zwahlen, C., Li, S.C., Kay, L.E., Pawson, T., and Forman-Kay, J.D. (2000). Multiple modes of peptide recognition by the PTB domain of the cell fate determinant Numb. EMBO J 19, 1505–1515. 10.1093/emboj/19.7.1505

41. Couturier, L., Mazouni, K., and Schweisguth, F. (2013). Numb localizes at endosomes and controls the endosomal sorting of notch after asymmetric division in Drosophila. Curr Biol 23, 588–593. 10.1016/j.cub.2013.03.002

42. Pylypenko, O., Lundmark, R., Rasmuson, E., Carlsson, S.R., and Rak, A. (2007). The PX-BAR membrane-remodeling unit of sorting nexin 9. EMBO J 26, 4788–4800. 10.1038/sj.emboj.7601889

43. Cabantous, S., Nguyen, H.B., Pedelacq, J.D., Koraichi, F., Chaudhary, A., Ganguly, K., Lockard, M.A., Favre, G., Terwilliger, T.C., and Waldo, G.S. (2013). A new protein-protein interaction sensor based on tripartite split-GFP association. Sci Rep 3, 2854. 10.1038/srep02854

44. Varnai, P., Lin, X., Lee, S.B., Tuymetova, G., Bondeva, T., Spat, A., Rhee, S.G., Hajnoczky, G., and Balla, T. (2002). Inositol lipid binding and membrane localization of isolated pleckstrin homology (PH) domains. Studies on the PH domains of phospholipase C delta 1 and p130. J Biol Chem 277, 27412–27422. 10.1074/jbc.M109672200

45. Rhee, S.G., and Bae, Y.S. (1997). Regulation of phosphoinositide-specific phospholipase C isozymes. J Biol Chem 272, 15045–15048. 10.1074/jbc.272.24.15045

46. Kim, J.H., Lee, C.-H., and Baek, M.-C. (2022). Dissecting exosome inhibitors: therapeutic insights into small-molecule chemicals against cancer. Experimental & Molecular Medicine 54, 1833–1843. 10.1038/s12276-022-00898-7

47. Catalano, M., and O’Driscoll, L. (2020). Inhibiting extracellular vesicles formation and release: a review of EV inhibitors. J Extracell Vesicles 9, 1703244. 10.1080/20013078.2019.1703244

48. Jeppesen, D.K., Fenix, A.M., Franklin, J.L., Higginbotham, J.N., Zhang, Q., Zimmerman, L.J., Liebler, D.C., Ping, J., Liu, Q., Evans, R., et al. (2019). Reassessment of Exosome Composition. Cell 177, 428–445 e418. 10.1016/j.cell.2019.02.029

49. Ratajczak, M.Z., and Ratajczak, J. (2020). Extracellular microvesicles/exosomes: discovery, disbelief, acceptance, and the future? Leukemia 34, 3126–3135. 10.1038/s41375-020-01041-z

50. Loughery, J., Cox, M., Smith, L.M., and Meek, D.W. (2014). Critical role for p53-serine 15 phosphorylation in stimulating transactivation at p53-responsive promoters. Nucleic Acids Res 42, 7666–7680. 10.1093/nar/gku501

51. Chen, X., Ko, L.J., Jayaraman, L., and Prives, C. (1996). p53 levels, functional domains, and DNA damage determine the extent of the apoptotic response of tumor cells. Genes Dev 10, 2438–2451. 10.1101/gad.10.19.2438

52. Marcellus, R.C., Teodoro, J.G., Charbonneau, R., Shore, G.C., and Branton, P.E. (1996). Expression of p53 in Saos-2 osteosarcoma cells induces apoptosis which can be inhibited by Bcl-2 or the adenovirus E1B-55 kDa protein. Cell Growth & Differentiation 7, 1643–1650.

53. Smith, C.A., Lau, K.M., Rahmani, Z., Dho, S.E., Brothers, G., She, Y.M., Berry, D.M., Bonneil, E., Thibault, P., Schweisguth, F., et al. (2007). aPKC-mediated phosphorylation regulates asymmetric membrane localization of the cell fate determinant Numb. Embo Journal 26, 468–480. 10.1038/sj.emboj.7601495

54. Chinnam, M., Xu, C., Lama, R., Zhang, X., Cedeno, C.D., Wang, Y., Stablewski, A.B., Goodrich, D.W., and Wang, X. (2022). MDM2 E3 ligase activity is essential for p53 regulation and cell cycle integrity. PLoS Genet 18, e1010171. 10.1371/journal.pgen.1010171

55. Wu, X., Bayle, J.H., Olson, D., and Levine, A.J. (1993). The p53-mdm-2 autoregulatory feedback loop. Genes Dev 7, 1126–1132. 10.1101/gad.7.7a.1126

56. van Neerven, S.M., and Vermeulen, L. (2023). Cell competition in development, homeostasis and cancer. Nat Rev Mol Cell Biol 24, 221–236. 10.1038/s41580-022-00538-y

57. Nichols, J., Lima, A., and Rodriguez, T.A. (2022). Cell competition and the regulative nature of early mammalian development. Cell Stem Cell 29, 1018–1030. 10.1016/j.stem.2022.06.003

58. Bowling, S., Lawlor, K., and Rodriguez, T.A. (2019). Cell competition: the winners and losers of fitness selection. Development 146. 10.1242/dev.167486

59. Baker, N.E., Kiparaki, M., and Khan, C. (2019). A potential link between p53, cell competition and ribosomopathy in mammals and in Drosophila. Dev Biol 446, 17–19. 10.1016/j.ydbio.2018.11.018

60. Bowling, S., Di Gregorio, A., Sancho, M., Pozzi, S., Aarts, M., Signore, M., M, D.S., Martinez-Barbera, J.P., Gil, J., and Rodriguez, T.A. (2018). P53 and mTOR signalling determine fitness selection through cell competition during early mouse embryonic development. Nat Commun 9, 1763. 10.1038/s41467-018-04167-y

61. Dejosez, M., Ura, H., Brandt, V.L., and Zwaka, T.P. (2013). Safeguards for cell cooperation in mouse embryogenesis shown by genome-wide cheater screen. Science 341, 1511–1514. 10.1126/science.1241628

62. Wagstaff, L., Goschorska, M., Kozyrska, K., Duclos, G., Kucinski, I., Chessel, A., Hampton-O’Neil, L., Bradshaw, C.R., Allen, G.E., Rawlins, E.L., et al. (2016). Mechanical cell competition kills cells via induction of lethal p53 levels. Nat Commun 7, 11373. 10.1038/ncomms11373

63. Zheng, C., Hu, Y., Sakurai, M., Pinzon-Arteaga, C.A., Li, J., Wei, Y., Okamura, D., Ravaux, B., Barlow, H.R., Yu, L., et al. (2021). Cell competition constitutes a barrier for interspecies chimerism. Nature 592, 272–276. 10.1038/s41586-021-03273-0

64. Venkatachalam, S., Shi, Y.P., Jones, S.N., Vogel, H., Bradley, A., Pinkel, D., and Donehower, L.A. (1998). Retention of wild-type p53 in tumors from p53 heterozygous mice: reduction of p53 dosage can promote cancer formation. EMBO J 17, 4657–4667. 10.1093/emboj/17.16.4657

65. Donehower, L.A., and Lozano, G. (2009). 20 years studying p53 functions in genetically engineered mice. Nat Rev Cancer 9, 831–841. 10.1038/nrc2731

66. Garcia-Cao, I., Garcia-Cao, M., Martin-Caballero, J., Criado, L.M., Klatt, P., Flores, J.M., Weill, J.C., Blasco, M.A., and Serrano, M. (2002). “Super p53” mice exhibit enhanced DNA damage response, are tumor resistant and age normally. EMBO J 21, 6225–6235. 10.1093/emboj/cdf595

67. Marruecos, L., Manils, J., Moreta, C., Gomez, D., Filgaira, I., Serafin, A., Canas, X., Espinosa, L., and Soler, C. (2021). Single loss of a Trp53 allele triggers an increased oxidative, DNA damage and cytokine inflammatory responses through deregulation of IkappaBalpha expression. Cell Death Dis 12, 359. 10.1038/s41419-021-03638-3

68. Hill, R., Song, Y., Cardiff, R.D., and Van Dyke, T. (2005). Selective evolution of stromal mesenchyme with p53 loss in response to epithelial tumorigenesis. Cell 123, 1001–1011. 10.1016/j.cell.2005.09.030

69. Ma, S., McGuire, M.H., Mangala, L.S., Lee, S., Stur, E., Hu, W., Bayraktar, E., Villar-Prados, A., Ivan, C., Wu, S.Y., et al. (2021). Gain-of-function p53 protein transferred via small extracellular vesicles promotes conversion of fibroblasts to a cancer-associated phenotype. Cell Rep 34, 108726. 10.1016/j.celrep.2021.108726

70. Wang, X., Song, K., Li, Y., Tang, L., and Deng, X. (2019). Single-Molecule Imaging and Computational Microscopy Approaches Clarify the Mechanism of the Dimerization and Membrane Interactions of Green Fluorescent Protein. Int J Mol Sci 20. 10.3390/ijms20061410

71. Ciferri, C., De Luca, J., Monzani, S., Ferrari, K.J., Ristic, D., Wyman, C., Stark, H., Kilmartin, J., Salmon, E.D., and Musacchio, A. (2005). Architecture of the human ndc80-hec1 complex, a critical constituent of the outer kinetochore. J Biol Chem 280, 29088–29095. 10.1074/jbc.M504070200

72. van der Wal, J., Habets, R., Varnai, P., Balla, T., and Jalink, K. (2001). Monitoring agonist-induced phospholipase C activation in live cells by fluorescence resonance energy transfer. J Biol Chem 276, 15337–15344. 10.1074/jbc.M007194200

73. Yarar, D., Surka, M.C., Leonard, M.C., and Schmid, S.L. (2008). SNX9 activities are regulated by multiple phosphoinositides through both PX and BAR domains. Traffic 9, 133–146. 10.1111/j.1600-0854.2007.00675.x

74. Cella Zanacchi, F., Lavagnino, Z., Perrone Donnorso, M., Del Bue, A., Furia, L., Faretta, M., and Diaspro, A. (2011). Live-cell 3D super-resolution imaging in thick biological samples. Nat Methods 8, 1047–1049. 10.1038/nmeth.1744

75. Sakuma, T., Nishikawa, A., Kume, S., Chayama, K., and Yamamoto, T. (2014). Multiplex genome engineering in human cells using all-in-one CRISPR/Cas9 vector system. Sci Rep 4, 5400. 10.1038/srep05400

76. Louis, N., Evelegh, C., and Graham, F.L. (1997). Cloning and sequencing of the cellular-viral junctions from the human adenovirus type 5 transformed 293 cell line. Virology 233, 423–429. 10.1006/viro.1997.8597

77. Berk, A.J. (2005). Recent lessons in gene expression, cell cycle control, and cell biology from adenovirus. Oncogene 24, 7673–7685. 10.1038/sj.onc.1209040

78. Godar, S., Ince, T.A., Bell, G.W., Feldser, D., Donaher, J.L., Bergh, J., Liu, A., Miu, K., Watnick, R.S., Reinhardt, F., et al. (2008). Growth-inhibitory and tumor-suppressive functions of p53 depend on its repression of CD44 expression. Cell 134, 62–73. 10.1016/j.cell.2008.06.006

79. Blattner, C., Hay, T., Meek, D.W., and Lane, D.P. (2002). Hypophosphorylation of Mdm2 augments p53 stability. Mol Cell Biol 22, 6170–6182. 10.1128/MCB.22.17.6170-6182.2002

80. Rappsilber, J., Ishihama, Y., and Mann, M. (2003). Stop and go extraction tips for matrix-assisted laser desorption/ionization, nanoelectrospray, and LC/MS sample pretreatment in proteomics. Anal Chem 75, 663–670. 10.1021/ac026117i

81. Matafora, V., Corno, A., Ciliberto, A., and Bachi, A. (2017). Missing Value Monitoring Enhances the Robustness in Proteomics Quantitation. J Proteome Res 16, 1719–1727. 10.1021/acs.jproteome.6b01056

82. Pelicci, S., Furia, L., Scanarini, M., Pelicci, P.G., Lanzano, L., and Faretta, M. (2022). Novel Tools to Measure Single Molecules Colocalization in Fluorescence Nanoscopy by Image Cross Correlation Spectroscopy. Nanomaterials (Basel) 12. 10.3390/nano12040686

83. Oneto, M., Scipioni, L., Sarmento, M.J., Cainero, I., Pelicci, S., Furia, L., Pelicci, P.G., Dellino, G.I., Bianchini, P., Faretta, M., et al. (2019). Nanoscale Distribution of Nuclear Sites by Super-Resolved Image Cross-Correlation Spectroscopy. Biophys J 117, 2054–2065. 10.1016/j.bpj.2019.10.036

84. Pelicci, S., Furia, L., Pelicci, P.G., and Faretta, M. (2023). Correlative Multi-Modal Microscopy: A Novel Pipeline for Optimizing Fluorescence Microscopy Resolutions in Biological Applications. Cells 12. 10.3390/cells12030354

85. Lobb, R.J., Becker, M., Wen, S.W., Wong, C.S., Wiegmans, A.P., Leimgruber, A., and Moller, A. (2015). Optimized exosome isolation protocol for cell culture supernatant and human plasma. J Extracell Vesicles 4, 27031. 10.3402/jev.v4.27031

86. Bosch, S., de Beaurepaire, L., Allard, M., Mosser, M., Heichette, C., Chretien, D., Jegou, D., and Bach, J.M. (2016). Trehalose prevents aggregation of exosomes and cryodamage. Sci Rep 6, 36162. 10.1038/srep36162

87. Bonsergent, E., Grisard, E., Buchrieser, J., Schwartz, O., Thery, C., and Lavieu, G. (2021). Quantitative characterization of extracellular vesicle uptake and content delivery within mammalian cells. Nat Commun 12, 1864. 10.1038/s41467-021-22126-y

