## Supplementary material for "Endocytic control of cell-autonomous and non-cell-autonomous functions of p53": merged file containing all supplementary figures

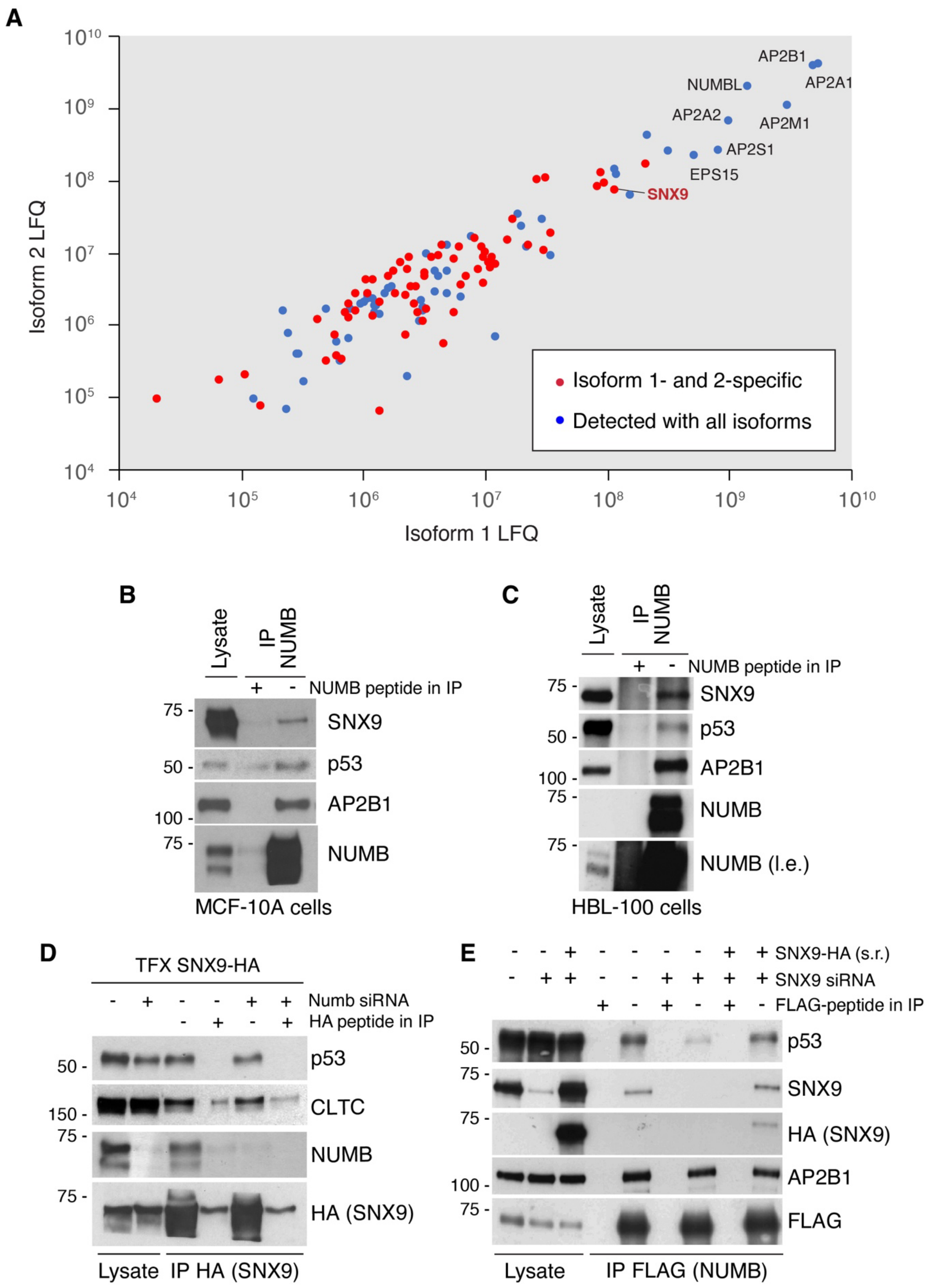


Supplementary Figure 1

**Figure S1. Additional data to Figure 1 of the main text. A.** Label-free quantification (LFQ) values of proteins identified in the NUMB-1 and NUMB-2 interactomes. NUMB-1/2–specific interactors are indicated in red, while those interacting with all isoforms are indicated in blue. Hits from the mass spec analysis were depleted of proteins present in the so-called "crapome" – a list of proteins frequently detected as non-specific contaminants in affinity purification mass spectrometry experiments [1] [http://www.crapome.org/]. The efficiency of the co-IP and mass spec analysis is supported by the detection of several known NUMB interactors (all isoforms), such as various AP2 adaptor complex subunits or EH-domain-containing proteins (EPS15, EPS15L1), indicated in the graph. **B,C.** MCF10A (**B**) or HBL-100 (**C**) cell lysates were IP with anti-NUMB and IB as indicated (right). A peptide corresponding to the NUMB epitope was used as negative control to compete antibody binding during the IP; l.e., long exposure. **D.** HEK-293 cells were silenced with NUMB siRNA (+) or Ctrl siRNA (-) and transfected with SNX9-HA as shown (top). Anti-HA IPs were IB as shown. An HA peptide was used as negative control to compete antibody binding during the IP. **E.** HEK-293 cells transfected with NUMB-1-FLAG were silenced with SNX9 siRNA (+) or Ctrl siRNA (-) and transfected with a siRNA-resistant SNX9-HA (s.r.) as shown (top). Anti-FLAG IPs were IB as shown (right). A FLAG peptide was used as negative control to compete antibody binding during the IP.

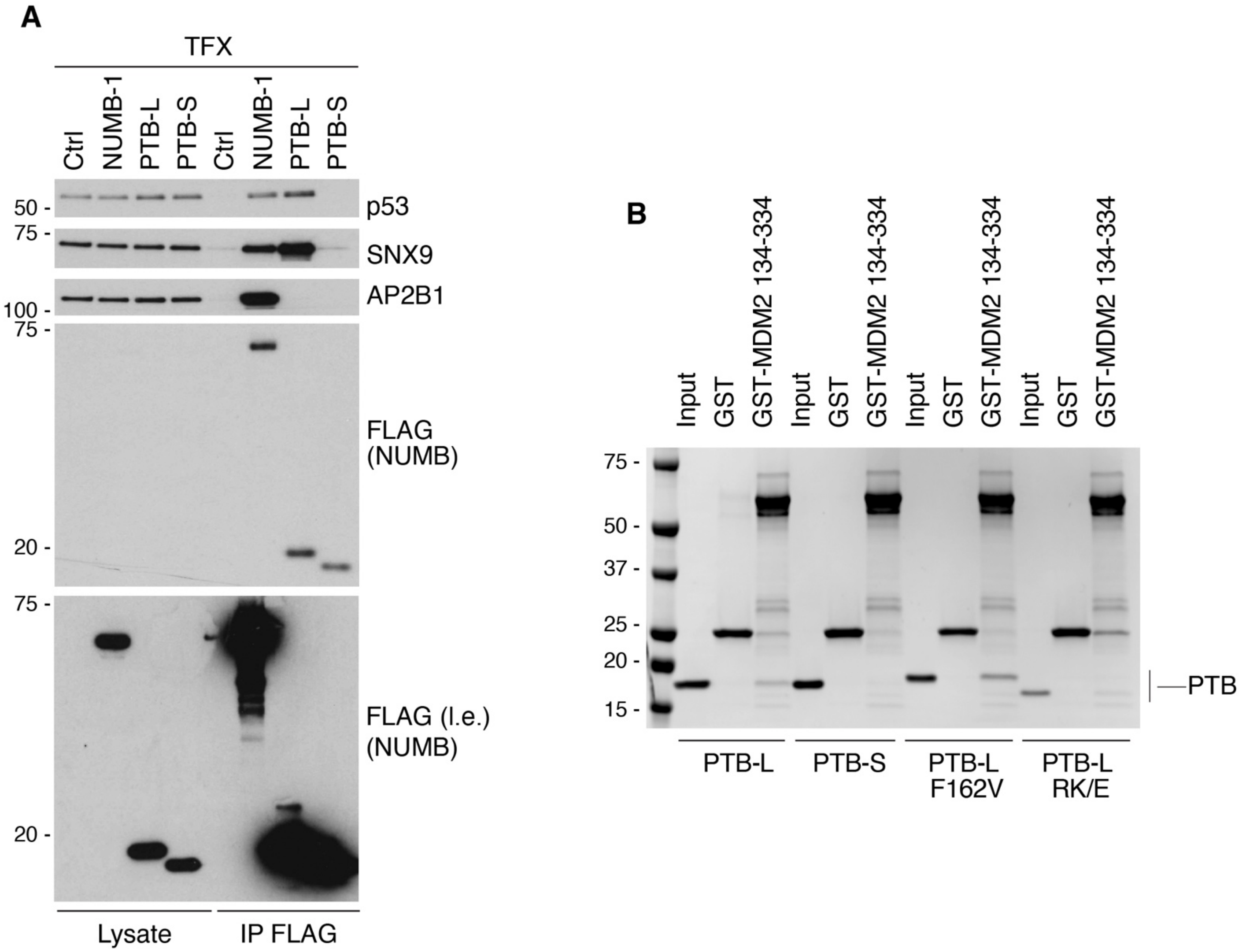


Supplementary Figure 2

**Figure S2. Additional data to Figures 2A and B of the main text. A.** The same blots as in Fig. 2A are shown, together with a long exposure (l.e.) of the anti-FLAG (NUMB) IB to allow visualization of the FLAG-tagged NUMB constructs in the total cellular lysate. **B.** *In vitro* binding assay performed with purified GST-MDM2 fragment 134-334 (containing the NUMB-interacting region [2]) or GST alone, immobilized on GSH beads, and the indicated purified NUMB-PTB proteins. Bound proteins were detected by Coomassie staining.

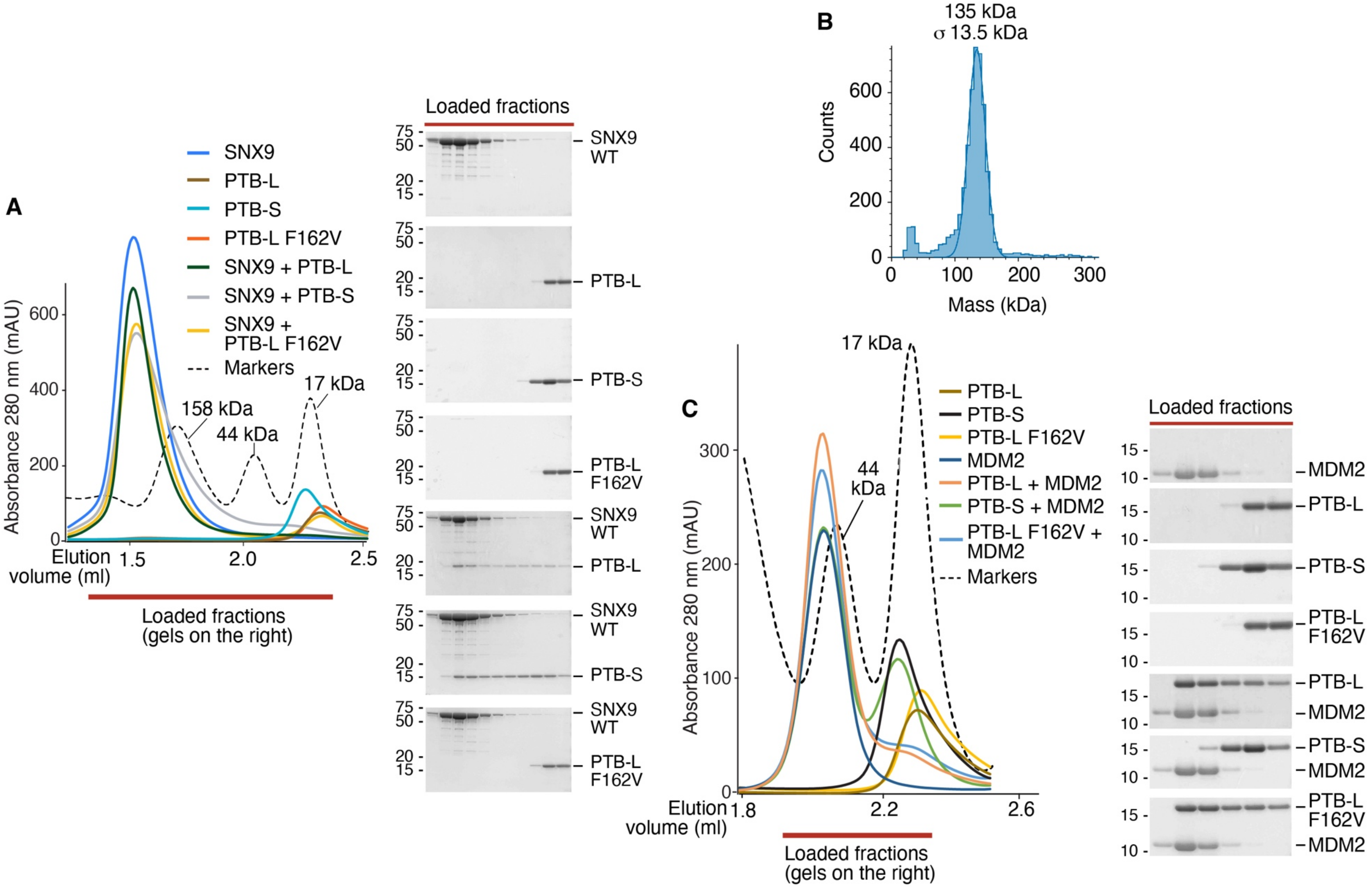


Supplementary Figure 3

**Figure S3. Additional data to Figure 2C and 2D of the main text. A.** Left, size-exclusion chromatography (SEC) elution profiles of the indicated purified proteins, alone or in combination. The red bar below the x-axis indicates the fractions loaded onto the gels shown on the right. Right, aliquots of the collected fractions were resolved by SDS-PAGE and proteins were detected by Coomassie staining. Note that the shift towards the left of PTB-L and PTB-S when combined with SNX9, compared with PTB-L and PTB-S alone, is indicative of the formation of a complex. This shift is not observed with the F162V mutant. Prior to SEC analysis, we observed minimal precipitation upon mixing certain species of SNX9 and PTB. This precipitate was removed by centrifugation of the sample before injection. This phenomenon may account for the slight reduction in peak absorbance observed for some SNX9/PTB complexes compared to SNX9 alone. The elution profile of the molecular weight marker shown here is consistent across all SEC experiments presented in the manuscript, as the same chromatography column was used throughout. **B.** As shown in panel A, SNX9 elutes in SEC at an apparent molecular weight above 158 kDa. To investigate this behavior, we analyzed purified full-length SNX9 by mass photometry. The main species detected had an average molecular mass of approximately 135 kDa, consistent with the formation of a SNX9 homodimer (monomer MW: 66.5 kDa), as expected given the dimerization properties of the SNX9 BAR domain [3]. σ 13.5 kDa represents the standard deviation. The slight discrepancy between the SEC elution profile and the calculated dimer mass may be attributed to the native conformation of SNX9, which can influence its hydrodynamic behavior. The y-axis of the graph shows the number of molecules analyzed (counts). **C.** SEC elution profiles of the indicated combinations of purified proteins (left). Fractions, indicated by the red bar, were analyzed SDS-PAGE and Coomassie staining (right). MDM2 corresponds to the MDM2 fragment aa 216-302, harboring phosphomimetic mutations which increase binding to NUMB [2]. The data confirm that the interaction between MDM2 and NUMB-PTB is largely dependent on the Ex3-encoded sequence of NUMB. Note that the $MDM2^{216\text{-}302}$ fragment alone eluted around the 44-kDa MW marker, as did the PTB-L/$MDM2^{216\text{-}302}$ complex. This is likely due to the intrinsically disordered nature of the $MDM2^{216\text{-}302}$ region, as previously reported [2], which causes the monomer to elute in SEC at an apparent molecular weight significantly higher than its theoretical mass. Some of the gels and SEC elution profiles of panel A are replicated in panel C (PTB proteins alone), because the two experiments were performed as a single experiment but are shown separately to avoid overcrowding of the tracks in the SEC elution profiles.

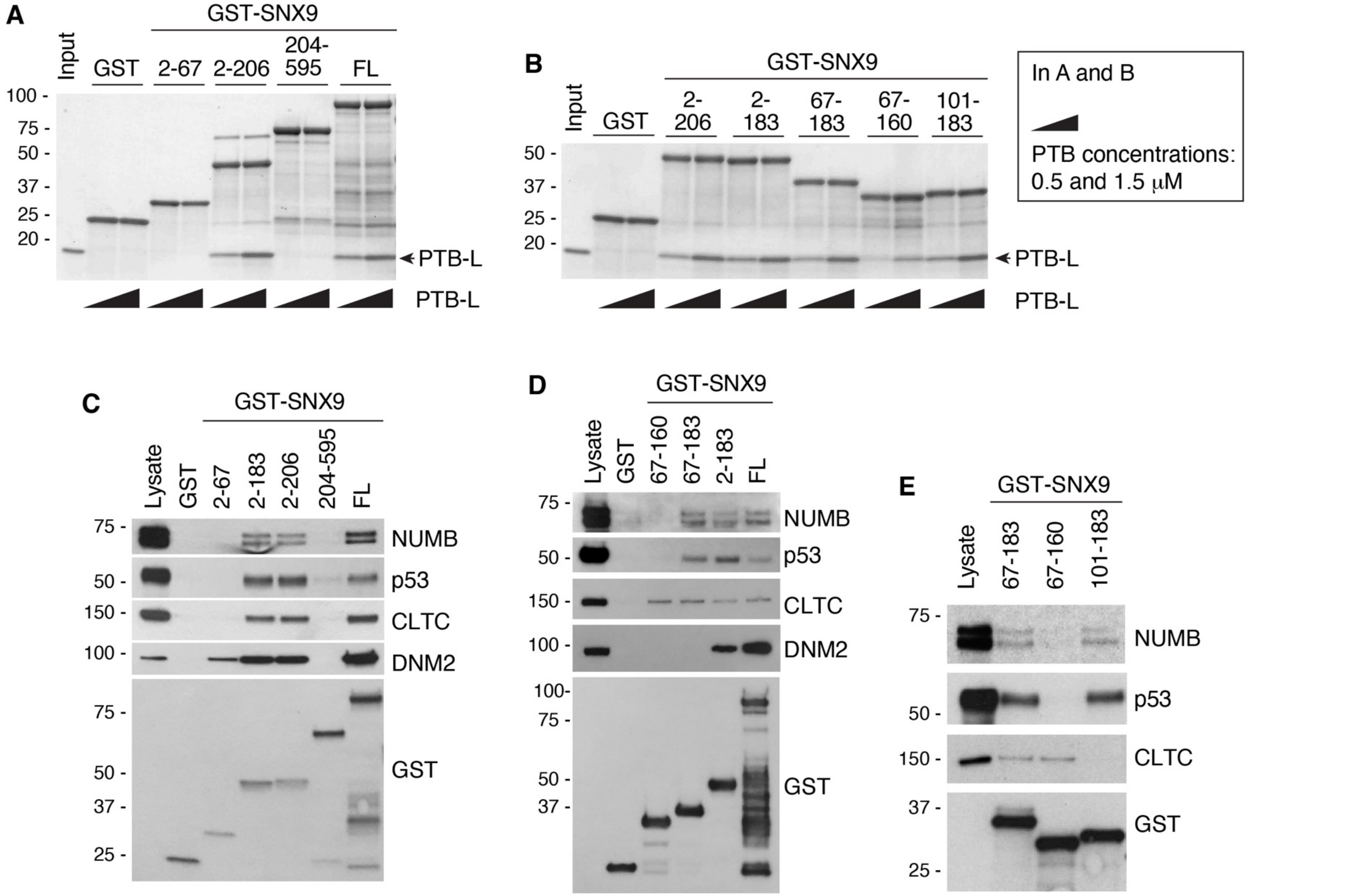


Supplementary Figure 4

**Figure S4. Additional data to Figure 2E of the main text: identification of the SNX9 LC domain as the binding surface for NUMB and p53. A, B.** *In vitro* binding assays with purified GST-SNX9 fragments or full-length (FL) protein immobilized onto GSH beads and increasing amounts of purified PTB-L (indicated by triangles). Bound proteins were detected by Coomassie staining. **C-E.** Pull-down assays performed with HEK-293 cell lysates and purified GST-SNX9 fragments or FL protein immobilized onto GSH beads, followed by IB as indicated on the right. DNM2 (dynamin 2) and CLTC (clathrin heavy chain) are known SNX9 interactors (see Fig. 2E), used here to confirm that the engineered SNX9 fragments display the expected binding abilities.

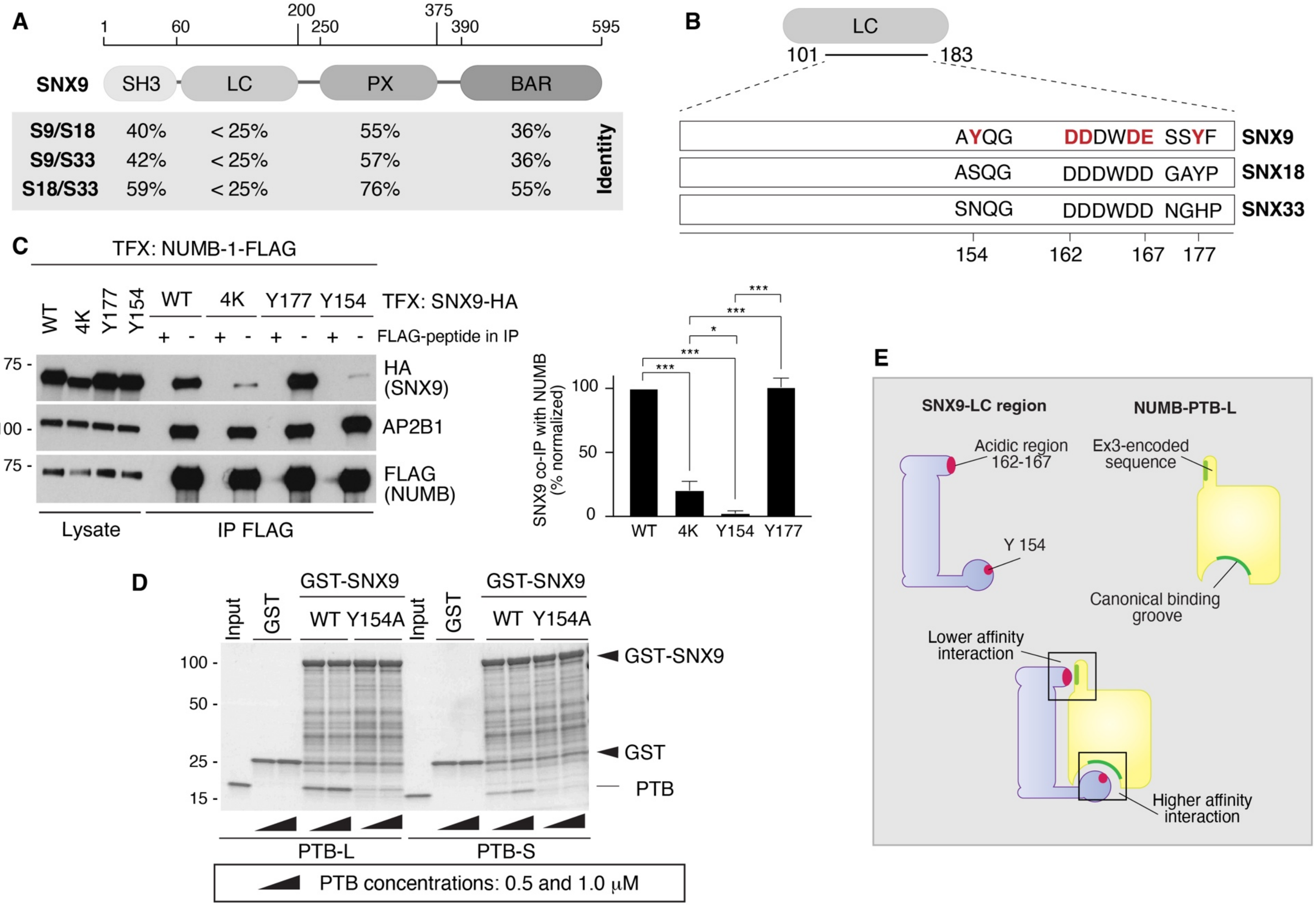


Supplementary Figure 5

**Figure S5. Additional data to Figure 2E of the main text: fine mapping of the SNX9 LC domain and deconvolution of the bidentate interaction with NUMB-PTB-L. A.** Domain organization of SNX9, SNX18 and SNX33 and sequence identities of the indicated domains, as per [4]. **B.** Alignment of the regions spanning residues 101 – 183 within the LC domains of SNX9, SNX18 and SNX33. Since SNX18 and SNX33 do not bind to NUMB-PTB-L (Fig. 1H), and since PTBs display preference for Y-based motifs for binding to their canonical binding groove [5], we reasoned that Y154 of SNX9, not present in SNX18 and SNX33, could be a candidate binding site. In addition, Y177 present in SNX9 and SNX18 but not in SNX33 could also be a candidate. To investigate the involvement of these Y residues (in red), we mutagenized them to A. The positions of the four acidic amino acids (D and E in red) mutagenized to K in the SNX9 4K mutant shown in panel **C** (see also main text) are also indicated. **C.** Left, HEK-293 cells were transfected (TFX) with NUMB-1-FLAG along with the indicated SNX9-HA constructs (WT or 4K, Y177 and Y154 mutants), followed by anti-FLAG IP and IB as shown (right). A FLAG peptide was used as negative control to compete with antibody binding during the IP Mut. 4K: SNX9 D162, D163, D166, E167>K. Mut. Y154: SNX9 Y154>A. Mut. Y177: SNX9 Y177>A. Right, quantitation of the amount of SNX9-HA (WT or mutant) binding to NUMB-1-FLAG by densitometry analysis of three independent co-IP experiments. The amount of SNX9 that co-IPs with NUMB was normalized to the level of expression of each SNX9-HA construct (see "lysate" lanes in the left panel) and expressed as a percentage of the co-IP between WT-SNX9 and NUMB. Results are presented as the mean ± SD. *, $P < 0.05$; ***, $P < 0.001$. **D.** *In vitro* binding assay performed with purified full-length GST-SNX9 wild type (WT) or mutant GST-SNX9-Y154A immobilized on GSH beads and increasing amounts (0.5 and 1.0 µM) of NUMB-PTB-L or NUMB-PTB-S (indicated by triangles). Bound proteins were detected by Coomassie staining. **E.** Model of the bidentate interaction between the SNX9 LC domain and NUMB PTB-L. The SNX9-Y154 residue mediates a higher affinity interaction with the PTB-binding groove. A lower affinity interaction is mediated by the acidic region of the SNX9 LC domain and the Ex3-encoded sequence of PTB-L.

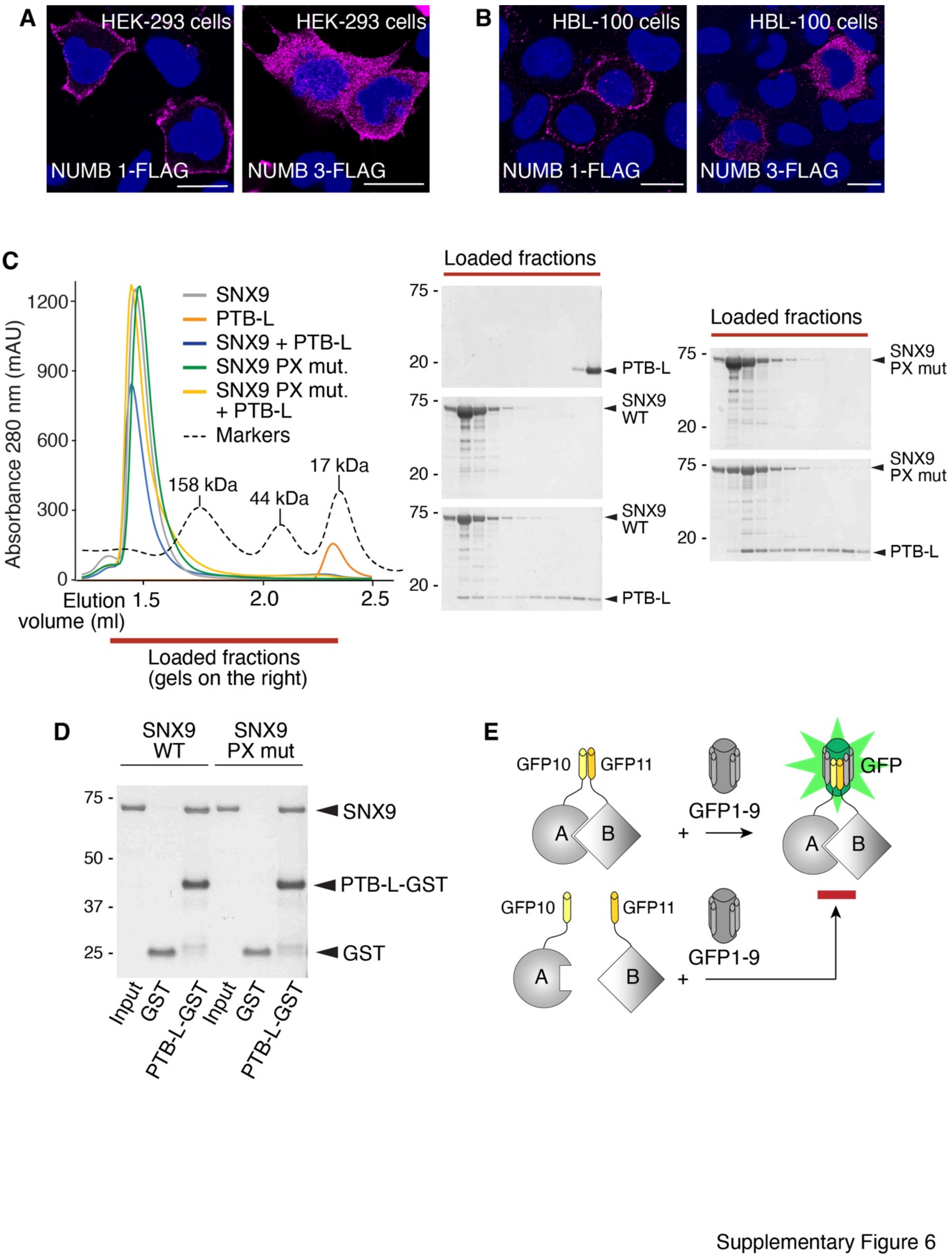


**Figure S6. Additional data to Figure 3 of the main text. A, B.** HEK-293 cells (**A**) or HBL-100 cells (**B**) were transfected with FLAG-tagged NUMB-1 or NUMB-3. Transfected NUMB was visualized by IF imaging with an anti-FLAG antibody (purple). Blue, DAPI counterstain. Bar, 20 µm. **C.** Size-exclusion chromatography (SEC) elution profiles of the indicated combinations of purified proteins (left). Fractions, indicated by the red bar, were analyzed by SDS-PAGE and Coomassie staining (right). **D.** *In vitro* binding assay with purified NUMB-PTB-L-GST or GST alone, immobilized on GSH beads, and purified SNX9-WT or SNX9-PXmut proteins. Bound proteins were detected by Coomassie staining. **E.** Scheme of the tripartite SPLIT-GFP system. Tripartite GFP complementation relies on the reconstitution of fluorescence through the interaction of three GFP fragments. Proteins of interest (A and B in the scheme) are tagged individually with two short fragments of GFP, GFP10 and GFP11. When A and B physically interact, GFP10 and GFP11 can self-assemble with a larger detector fragment, GFP1-9, expressed in the cell, enabling proper GFP folding and chromophore maturation (top). The resulting fluorescence allows the precise detection of protein-protein interactions or colocalization within the cell with minimal background signal. When A and B fail to interact the GFP fluorescence signal is not emitted (bottom).

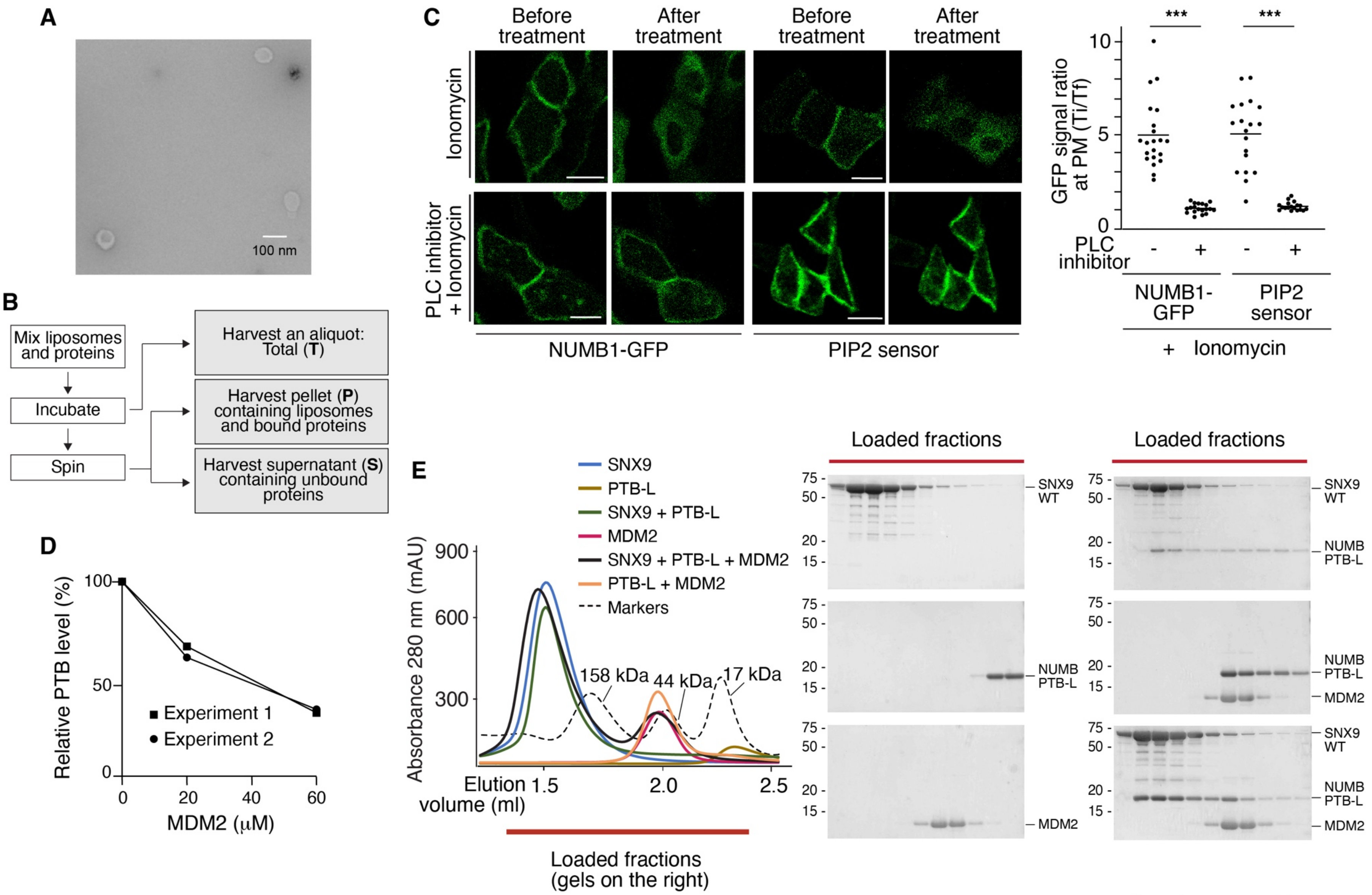


Supplementary Figure 7

**Figure S7. Additional data to Figure 4 of the main text. A.** Representative transmission electron microscopy images of liposomes used in the liposome binding assays described in the main text. Bar, 100 nm. **B.** Scheme of the liposome binding assay. **C.** MCF10A cells were transfected with NUMB1-GFP or a PIP2 sensor-YFP (pleckstrin homology domain of PLCδ1 fused to YFP) and treated as indicated. Where indicated, cells were treated with the PLC inhibitor U73122 (10 μM) or DMSO control for 3 min before adding ionomycin (10 μM) for 6 min. Left. Representative GFP or YFP epifluorescence (green) images from time-lapse microscopy before and after treatment. The fluorescence signal is displayed using the same color scale for all samples, as images were acquired using identical excitation and emission wavelengths compatible with both fluorophores. Bar 20 μm. Right. Quantitation of the experiment. For each cell analyzed, the green signal at the PM was quantified at the beginning (Ti) and end of treatment (Tf). The average Ti/Tf ratio for each analyzed cell was plotted as an individual data point. Results are from two independent experiments, with the mean for each condition indicated by the horizontal line. ***, $p < 0.001$. **D.** Quantitation of the experiment in Fig. 4B, related to the PTB-L/$MDM2^{216\text{-}302}$ combination, repeated on two independent experiments. **E.** Left, size-exclusion chromatography (SEC) elution profiles of the indicated combinations of purified proteins. Fractions, indicated by the red bar were analyzed by SDS-PAGE and Coomassie staining (right). Note that the PTB-L profiles shifts to the left when combined with SNX9 or $MDM2^{(216\text{-}302)}$ compared with PTB-L alone, indicating the formation of dimeric complexes. When all three proteins are present, a tripartite complex is not formed; rather PTB-L forms independent dimeric complexes with SNX9 and $MDM2^{(216\text{-}302)}$. This finding is supported by the fact that $MDM2^{(216\text{-}302)}$ is absent in the fractions enriched in the PTB-L:SNX9 complex. The gels and SEC elution profiles (with the exception of the combination PTB-L:SNX9:MDM2) are replicated from Figs. S3A,C because the experiments were performed as a single experiment but are shown separately to avoid overcrowding of the tracks in the SEC elution profiles.

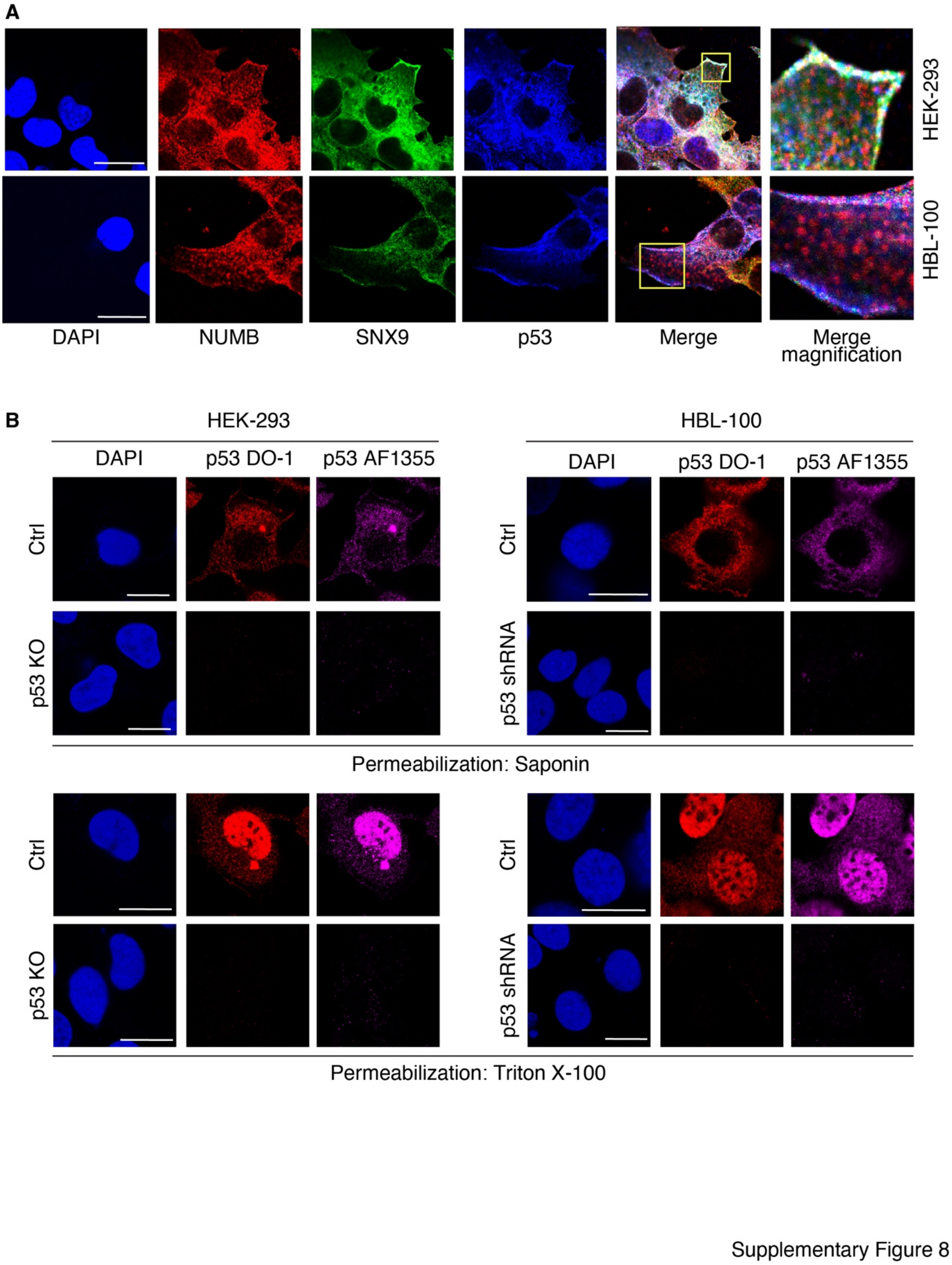


**Figure S8. Additional data to Figure 5 of the main text. A.** IF analysis of HEK-293 cells (top) and HBL-100 cells (bottom) with anti-NUMB (red), anti-SNX9 (green), and anti-p53 (blue) antibodies. Cell permeabilization was performed with saponin to permeabilize preferentially the PM, allowing for a clearer assessment of cytoplasmic and PM localized p53 by minimizing the nuclear signal. This approach was necessary due to the significantly higher levels of nuclear p53 compared to cytoplasmic levels. Merge with magnification of boxed areas is shown on the right. DAPI, nuclear stain. Bar, 20 µm. **B.** The panel shows a series of controls to illustrate the specificity of the staining obtained with the two anti-p53 Abs used in the study (DO-1 and AF1355, stained as indicated on top) and the different effects of solubilization with saponin or Triton X-100 (upper panels and lower panels). Specificity was shown in HEK-293 (left panels) and HBL-100 (right panels) cells by ablating p53 expression either by KO (in HEK-293) of by shRNA (in HBL-100). As shown, the permeabilization with saponin, which leaves the nuclear membrane largely intact, allows for better appreciation of the non-nuclear pool of p53, which is predominant in the cell, as shown by permeabilizing the cells with Triton X-100. DAPI, nuclear counterstain. Bar, 20 µm.

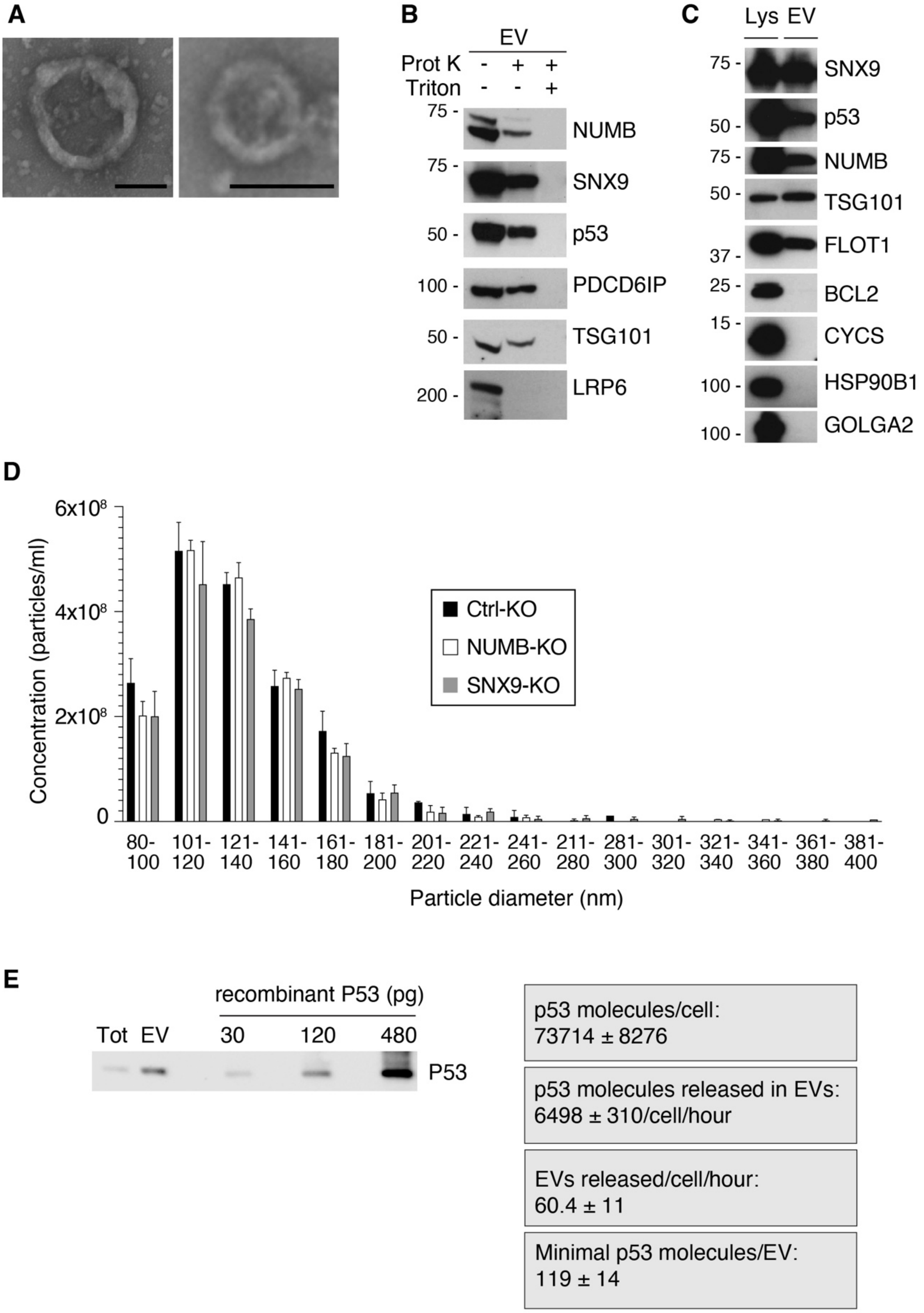


Supplementary Figure 9

**Figure S9. Additional data to Figure 6 of the main text. A**. The quality of the EV purification was controlled by visualizing vesicles using electron transmission microscopy. Representative images of EVs purified from the conditioned medium of HEK-293 cells are shown; bar 100 nm. **B.** To demonstrate that proteins present in EVs preparations are protected by a phospholipid bilayer, we treated them with Proteinase K (Prot K). EVs purified as in 'A' were treated with Prot K (5 μg/ml) and Triton X-100 (1% final) as indicated and IB as shown on the right. **The partial decrease in the protein levels in EVs following Prot K treatment** may be caused by multiple factors, as previously reported [6,7]: (i) the high shear stress generated during resuspension of the ultracentrifugation pellet could result in partial vesicle disruption; and (ii) cleavage of membrane-associated EV proteins might compromise vesicle integrity, leading to increased susceptibility to proteolytic degradation. **C.** In the experiments shown in Fig. 6A, 6C, 6G, S9A, S9B, S9D, we used EVs purified from HEK-293 cells. In the experiments shown in Fig. 6B, 7C, 7D, and S10C, we used EVs purified from Expi293 cells, a derivative of HEK-293 cells that grows in suspension at high density, thus allowing a higher yield of purified EVs. We demonstrated that EVs derived from Expi293 cells show the same characteristics as those derived from HEK-293 cells, especially in terms of the amount of SNX9, NUMB, p53 and other markers packaged into EVs, as shown by IB (compare with Fig. 6A). Lys, total cellular lysate. **D.** EVs from HEK-293 Ctrl-KO, SNX9-KO and NUMB-KO cells, as described in Fig. 6G, were analyzed by tunable resistive pulse sensing (TRPS) to measure concentration and diameter. Results are expressed as means ± SD of three technical replicates. The one-way ANOVA test was not significant, indicating that the KO of SNX9 or NUMB does not affect the number or size of EVs. **E.** MCF10A cells were cultivated to near confluence (12 x $10^6$ cells/15 cm plate), washed three times with PBS, and re-incubated in serum-free medium for 20 min. The EVs released in this 20 min window were purified from the conditioned medium. Purified EVs (1/13$^{th}$ of total preparation) was analyzed IB as indicated (EV lane), alongside total cellular lysate (1/2380$^{th}$ of total lysate) and the indicated amounts of purified recombinant p53. Densitometry analysis of two independent experiments was performed to quantify the amount of p53 present in cells and EVs. Right, the number of p53 molecules/cell, number of p53 molecules released in EVs/cell/h, number of EVs produced by a cell/hour, and the minimal number of p53 molecules present in a single EV are shown (mean ± SD).

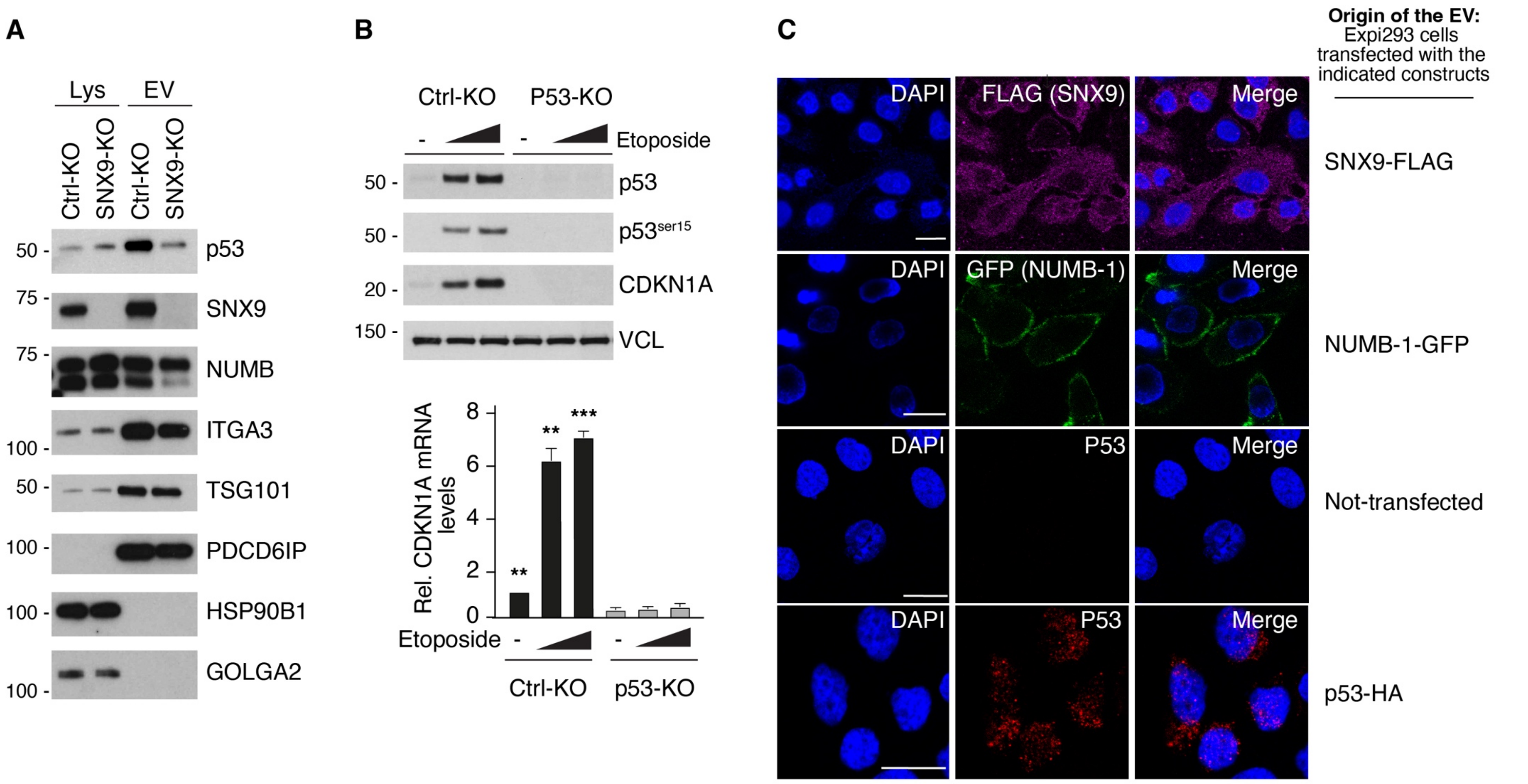


Supplementary Figure 10 A-C

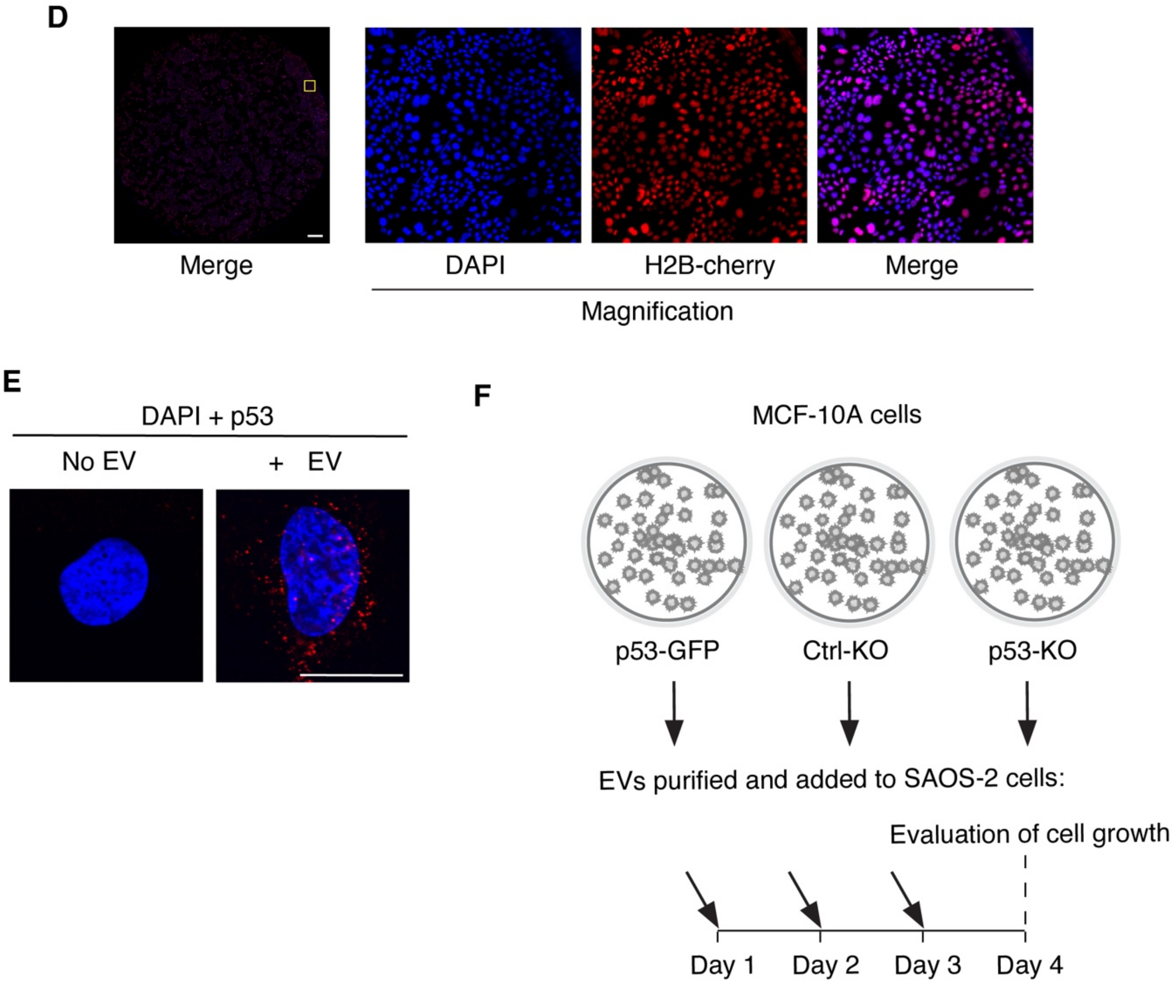


Supplementary Figure 10 D-F

**Figure S10. Additional data to Figure 7 of the main text. A.** EVs were purified from the conditioned medium of MCF10A Ctrl-KO or SNX9-KO cells and the levels of p53, SNX9 and NUMB were analyzed by IB. ITGA3 (integrin α3), TSG101 and PDCD6IP (Alix) are positive EVs markers. HSP90B1 and GOLGA2 are negative EVs markers. Lys, total cellular lysate. **B.** MCF10A cells Ctrl-KO or p53-KO MCF10A cells were treated with increasing doses of etoposide (as in Fig. 7A, B) and analyzed by IB (top) or RT-qPCR (bottom). For the RT-qPCR analysis, data are from three independent experiments and expressed as mean ± SD. ** and ***, $p < 0.01$ and $< 0.001$, respectively, *vs*. same condition in p53 KO. Vinculin (VCL) is used as a loading control. **C.** Expi293 cells were transfected as indicated on the right (SNX9-FLAG, NUMB-1-GFP, not-transfected, p53-HA) and EVs were purified from their conditioned medium. EVs were then added to MCF10A-p53-KO recipient cells for 8 h. Recipient cells treated with p53-HA EVs were also treated with etoposide (50 µM) for 8 h. IF of recipient cells was performed as indicated to visualize the tagged constructs (SNX9-FLAG, purple; NUMB-GFP, green; p53-HA, red). Blue, DAPI counterstain. Bar 20 µm. **D**. The coverslips harvested as per point d in Fig. 7E were analyzed for possible contamination of cells from the co-culture plate. Coverslips were counterstained with DAPI and analyzed for coincidence of the DAPI and H2B-Cherry signals. The leftmost panel shows an entire coverslip; two coverslips for a total of 100,553 cells were acquired using a HC PL FLUOTAR 10X objective mounted on a DMi8 inverted microscope (Thunder Imaging System - Leica Microsystems). The yellow inset refers to the region of the coverslip magnified in the other three panels. In the two analyzed coverslips, the % of overlap between the DAPI and H2B-Cherry signals was 99.8 and 99.3%. We concluded that virtually no cross-contamination of the coverslips by the co-cultivated cells occurred. Bar 1 mm. **E**. SAOS2 cells were treated for 15 h with EVs purified from the conditioned medium of MCF10A-p53-GFP cells (+ EV) or left untreated (No EV). Cells were also treated with the proteasome inhibitor MG132 (5 μM) to stabilize p53 and facilitate its detection. IF was performed with anti-p53 (red) and DAPI (blue). Bar, 20 μm. **F**. Scheme of the experiment of Fig. 7G. EVs were purified from the conditioned media of MCF10A-Ctrl-KO, MCF10A-p53-KO, and MCF10A-p53-GFP cells, and added to SAOS2 cells for three days. Cell growth was then evaluated on day 4.

### REFERENCES

1. Mellacheruvu, D., Wright, Z., Couzens, A.L., Lambert, J.P., St-Denis, N.A., Li, T., Miteva, Y.V., Hauri, S., Sardiu, M.E., Low, T.Y., et al. (2013). The CRAPome: a contaminant repository for affinity purification-mass spectrometry data. Nat Methods 10, 730-736. https://doi.org/10.1038/nmeth.2557
2. Colaluca, I.N., Basile, A., Freiburger, L., D'Uva, V., Disalvatore, D., Vecchi, M., Confalonieri, S., Tosoni, D., Cecatiello, V., Malabarba, M.G., et al. (2018). A Numb-Mdm2 fuzzy complex reveals an isoform-specific involvement of Numb in breast cancer. J Cell Biol 217, 745-762. https://doi.org/10.1083/jcb.201709092
3. Pylypenko, O., Lundmark, R., Rasmuson, E., Carlsson, S.R., and Rak, A. (2007). The PX-BAR membrane-remodeling unit of sorting nexin 9. EMBO J 26, 4788-4800. https://doi.org/10.1038/sj.emboj.7601889
4. Haberg, K., Lundmark, R., and Carlsson, S.R. (2008). SNX18 is an SNX9 paralog that acts as a membrane tubulator in AP-1-positive endosomal trafficking. J Cell Sci 121, 1495-1505. https://doi.org/10.1242/jcs.028530
5. Zwahlen, C., Li, S.C., Kay, L.E., Pawson, T., and Forman-Kay, J.D. (2000). Multiple modes of peptide recognition by the PTB domain of the cell fate determinant Numb. EMBO J 19, 1505-1515. https://doi.org/10.1093/emboj/19.7.1505
6. Bonsergent, E., Grisard, E., Buchrieser, J., Schwartz, O., Thery, C., and Lavieu, G. (2021). Quantitative characterization of extracellular vesicle uptake and content delivery within mammalian cells. Nat Commun 12, 1864. https://doi.org/10.1038/s41467-021-22126-y
7. Foers, A.D., Chatfield, S., Dagley, L.F., Scicluna, B.J., Webb, A.I., Cheng, L., Hill, A.F., Wicks, I.P., and Pang, K.C. (2018). Enrichment of extracellular vesicles from human synovial fluid using size exclusion chromatography. J Extracell Vesicles 7, 1490145. https://doi.org/10.1080/20013078.2018.1490145
